# Synthesis and characterization of novel intrinsically fluorescent analogs of cholesterol with improved photophysical properties

**DOI:** 10.1101/2023.11.23.568457

**Authors:** Max Lehmann, Peter Reinholdt, Mohammad Bashawat, Holger A. Scheidt, Senjuti Halder, Duccio di Prima, Jacob Kongsted, Peter Müller, Pablo Wessig, Daniel Wüstner

## Abstract

Live-cell imaging of cholesterol trafficking depends on suitable cholesterol analogs. However, existing fluorescent analogs of cholesterol either show very different physico-chemical properties compared to cholesterol or demand excitation in the ultraviolet spectral region. We present novel intrinsically fluorescent sterols containing four conjugated double bonds in the ring system and either a hydroxy or a keto group in the C3 position. Synthesis of these probes involves dehydrogenation of 7-dehydrocholesterol using mercury(II) acetate, Swern oxidation/dehydrogenation, and stereoselective Luche reduction. Molecular dynamics simulations and nuclear magnetic resonance spectroscopy reveal that the analog with a 3’-hydroxy-group like cholesterol can condense fatty acyl chains and form hydrogen bonds to water molecules at the bilayer interface. The emission of both probes is red-shifted by 80-120 nm compared to the widely used sterol analogs dehydroergosterol or cholestatrienol. This allows for their imaging on conventional microscopes, as we here show in giant unilamellar vesicles. These experiments reveal a preferred partitioning of both sterol probes into the biologically relevant liquid-ordered phase. In conclusion, we present a synthesis strategy leading to novel intrinsically fluorescent sterol probes with close resemblance of cholesterol. Their improved photophysical properties will allow for live-cell imaging of sterol transport in the future.

## Introduction

Cholesterol is the most abundant lipid species in the plasma membrane (PM) of mammalian cells, playing a central role in its function by regulating membrane packing, protein dynamics, permeability, and membrane trafficking.^[1]^ Biophysical and cell biological studies have shown that cholesterol possesses unique properties enabling this sterol to fine-tune membrane properties.^[1a, 1b]^ Trafficking of cholesterol between subcellular membranes is an important aspect/process of cellular cholesterol homeostasis. Disturbed cholesterol transport is observed in a variety of human diseases, ranging from atherosclerosis and lysosomal storage disorders over neurodegenerative diseases to cancer.^[1a, 2]^ Studying intracellular cholesterol transport heavily relies on fluorescence microscopy, and several approaches have been developed to visualize cholesterol in cells. One approach is based on using fluorescent polyene macrolides, such as filipin, binding to cholesterol and other sterols with free 3’-hydroxy groups, which contains intrinsic fluorescence due to conjugated double bonds in the polyene portion of the molecule, allowing for their visualization by fluorescence microscopy. Filipin can be conveniently visualized using DAPI filters available on most wide field and confocal microscopes, which explains its widespread use. Significant disadvantages of this polyene, however, are, that (i) cells need to be fixed for filipin staining, and (ii) that no particular sterol pool can be directly monitored (e.g., newly synthesized versus sterols internalized by cells).^[3]^ Also, filipin staining cannot distinguish cholesterol from its biosynthetic precursors nor from oxysterols. An alternative approach is to use cholesterol binding toxins, such as as perfringolysin O (PFO) or aerolysin D4 tagged with a suitable fluorescent reporter.^[4]^ Such toxins bind to cholesterol-rich model and cell membranes, but only above a certain threshold concentration, often defined as ‘accessible cholesterol’. The nature of this pool is not clear, and suitability of such toxins for quantifying membrane cholesterol has been debated.^[4c, 5]^ A third strategy to follow cholesterol trafficking by microscopy is to use fluorescent cholesterol analogs bearing a covalently linked dye, preferentially attached to the side chain of cholesterol. Here, BODIPY- or NBD-moieties are often employed, allowing for convenient live-cell imaging of sterol transport and even for correlative and super-resolution microscopy.^[6]^ A disadvantage of these probes is their limited resemblance of cholesterol, since the attached dye significantly affects the biophysical properties of such analogs, leading e.g. to strongly reduced ability to condense membranes or even to opposite membrane orientation compared to cholesterol.^[7]^ A variant of this approach is to use alkyne derivatives of cholesterol to which a fluorescent moiety can be attached after cells have been labeled using bioorthogonal click chemistry.^[6a, 8]^ While the alkyne analogs are only minimally modified variants of cholesterol with very similar properties, the click reaction and the resulting dye-attached cholesterol analog can disturb membrane properties and transport in hardly controllable ways.

Fourth, cholesterol analogs have been synthesized and employed which contain conjugated double bonds endowing the molecules with an intrinsic fluorescence. Cholesterol contains a single Δ5 double bond in the ring system, while the yeast sterol ergosterol contains additionally a Δ6 double bond as well as an extra methyl group and double bond in the side chain. Intrinsically fluorescent sterols, such as the cholesterol analog cholestatrienol (CTL) or the ergosterol analog dehydroergosterol (DHE), contain three conjugated double bonds in the ring system including the natural ones, giving such probes a slight fluorescence in the ultraviolet (UV) region of the spectrum, while maintaining very similar biophysical and biochemical properties like cholesterol and ergosterol, respectively.^[7a, 9]^ These analogs can replace cholesterol and ergosterol in sterol auxotroph organisms, interact with sterol binding proteins and can be used in spectroscopic and imaging assays.^[10]^ Another advantage is that the same strategy of introducing additional double bonds into the ring system can be employed to generate analogs of important bioactive cholesterol derivatives, such as oxysterols or steroid hormones.^[11]^ Disadvantages of such sterol molecules like DHE or CTL are that their visualization requires UV-optimized microscope optics, and that they have low brightness and high bleaching propensity making long-term imaging impossible.^[6b, 12]^ Few studies have attempted to use multi-photon excitation for imaging of DHE and CTL, but here, the low absorption cross-section demands long acquisition times, thereby preventing routine applications.^[9b, 13]^ This is particularly pronounced in cellular studies, where three-photon excitation was used to minimize parallel excitation of cellular autofluorescence.^[9b, 10d, 13a]^

A strategy to overcome such limitations is to extend the conjugated system by introducing a fourth double bond into the ring system. Computational studies predict, that this modification should result in strongly improved photophysical properties with pronounced red-shift in excitation and emission, higher oscillator strength and increased multi-photon absorption cross section.^[11b, 14]^ A first experimental realization of this strategy was performed for analogs of cholesterol and ergosterol with four double bonds in the ring system and provided promising results by confirming red-shifted emission which allowed for imaging of these probes using conventional wide field optics.^[15]^ Extension of the conjugated system also resulted in much reduced photobleaching and thereby in prolonged live-cell imaging of these probes. Due to rapid keto-enol tautomerization, however, these analogs are converted into the corresponding polyenes with three double bonds and a 3’-keto group, necessitating the protection of the 3’-hydroxy moiety with an acetate group.^[15]^ To overcome this limitation, the positions of the conjugated double bonds would need to be reallocated, while maintaining the biophysical properties of the natural sterols. This could prevent the keto-enol tautomerization and theoretically also affect the photophysical properties of such sterol probes while maintaining the biophysical properties, as predicted by electronic structure calculations and molecular dynamics (MD) simulations, respectively.^[14]^

Here, we present two new intrinsically fluorescent sterol analogs, each with four double bonds distributed over the A-, B- and C-ring of the steroid ring system and containing the alkyl chain of cholesterol. These probes have an identical core structure but differ in the 3’-carbon position, where **5** contains a keto group, while **6** has a hydroxy group, like cholesterol. The photo-physical and membrane properties of the molecules were characterized using a couple of experimental and theoretical approaches. Both sterol probes have a pronounced red-shift in their excitation and emission, which amounts to about 80 nm compared to DHE and CTL. Together with a much-improved photostability this allows for their convenient imaging on conventional wide field and confocal microscopes. We find that **5** and **6** are further red-shifted in polar and protic compared to apolar solvents whereby **5** has a fluorescence intensity approx. 10 times higher than **6**. Both probes can be incorporated into lipid membranes and detected using a conventional wide field microscope. Notably, **6** is able to order fatty acyl chains of phospholipids, and both sterol probes partition into the liquid ordered phase of giant unilamellar vesicles (GUVs), thereby resembling the properties of endogenous cholesterol.

Together, we present novel fluorescent probes which at the same time resemble the respective physiological sterols closely and exhibit improved optical properties allowing for their quantitative imaging on conventional microscope systems in future cellular studies.

## Results and Discussion

### Synthesis of 5 and 6

The synthetic route to **5** and **6** starts from commercially available 7-dehydrocholesterol **1** (7-DHC, vitamin-D_3_ precursor). After protection of the hydroxy group as acetate (**2**), the third C-C-double bond is installed upon treatment with mercury(II) acetate. This kind of dehydrogenation of steroids was discovered nearly a century ago by Windaus and Linsert with the conversion of ergosterol to DHE^[16]^ and later applied to the synthesis of cholestatrienol (CTL) **4** from **1**^[17]^ and cholestatrienol acetate **3** from **2**.^[18]^ Compared with these works, we could considerably increase the yield of **3**. The subsequent saponification of **3** to **4** proceeds with nearly quantitative yield (Scheme 1).

Wilson and co-workers report in 2000 the unintentional formation of tetraenone **5** under Swern oxidation^[19]^ conditions using a large excess of reagents.^[20]^ “Over-oxidation” with formation of α,β-unsaturated ketones was occasionally observed as side effect of the Swern oxidation.^[21]^ It might be reasonable to assume that the Swern intermediates (Me_2_S-Cl)^+^ are responsible for this outcome, which may act as Cl^+^ sources.^[22]^ A plausible mechanism for the formation of **4** from **5** is summarized in Scheme 2. After “normal” Swern oxidation of 3’-OH group the resulting ketone **A** is deprotonated to give the highly resonance-stabilized enolate **B**. The chlorination of **B** with (Me_2_S-Cl)^+^ affords the intermediate **C**. In the final step to **5** an elimination of HCl occurs in the presence of the base NEt_3_ (Scheme 2).

The tetraenol **6** is obtained by stereoselective reduction of the keto group in **5** under Luche conditions (CeCl_3_/NaBH_4_).^[23]^ The 3β-configuration in **6** was unambiguously proven by NMR-spectroscopy (Scheme 1, for details see the Supporting Information).

**Scheme 1.**
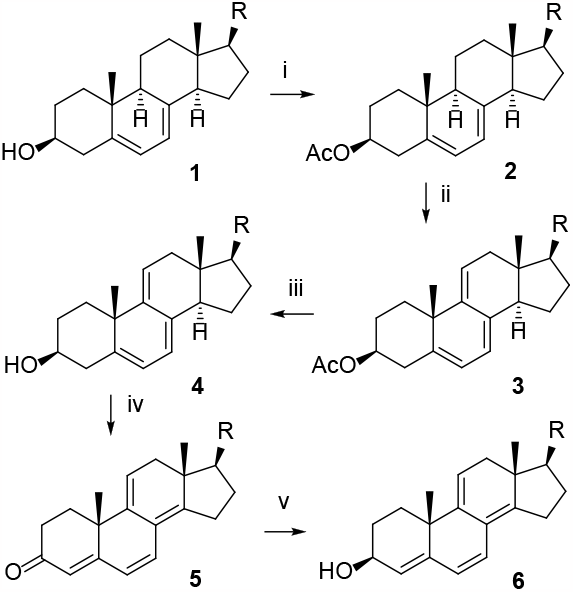
Synthesis of tetraenone **5** and tetraenol **6** (i: AcCl, Py, 0 °C -> r.t., 24h, 100%. ii: Hg(OAc)_2_, AcOH, EtOH/DCM, 48h, r.t., 65%. iii: NaOH, NeOH, rfx. 1h, 98%. iv: 10eq. (COCl)_2_, 20 eq. DMSO, 10 eq. NEt_3_, -76°C, 78%. v: CeCl_3_(H_2_O)_7_, 0°C, 60 min, 55%, R =C_8_H_17_)

**Scheme 2.**
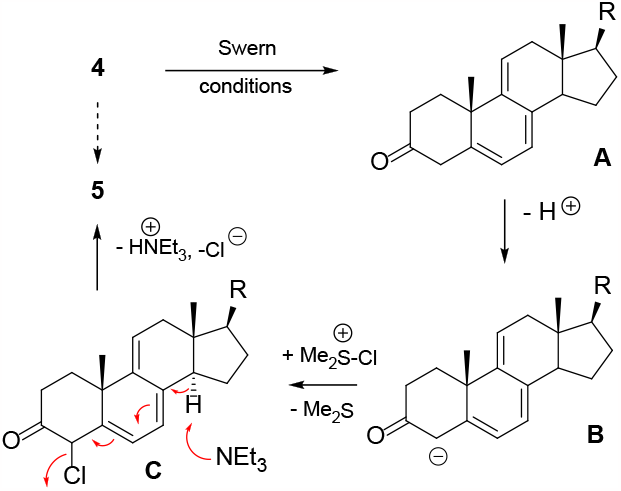
Mechanistic proposal for the formation of tetraenone **5** (R = C_8_H_17_).

### Optical properties of 5 and 6

The photophysical properties of the fluorescent sterol probes were first studied using electronic structure calculations. Table 1 reports the calculated excitation energies and one-, two- and three-photon transition strengths for the fluorescent analogs of cholesterol. **5** has two low-lying states: a near-forbidden n-π* state, and a π-π* state, which is bright in a one-photon absorption process. The absorption energy is predicted to be redshifted compared to that of DHE and CTL by about 0.3 eV (Table 1). The two states are very close in energy (around 0.16 eV), which means that the ordering of the two states can likely change based on environmental factors such as the polarity of the surrounding medium. Such a phenomenon has been observed previously for 3’-ketosteroids with conjugated double bonds.^[24]^

**Table 1.** Comparison of transition energies and oscillator strengths of **5** and **6** versus DHE and CTL.

| | CTL<br>$\pi-\pi^*$<br>S1 | DHE<br>$\pi-\pi^*$<br>S1 | **5**<br>$n-\pi^*$<br>S1 | **5**<br>$\pi-\pi^*$<br>S2 | **6**<br>$\pi-\pi^*$<br>S1 | **6**<br>$\pi-\pi^*$<br>S2 |
| --- | --- | --- | --- | --- | --- | --- |
| $\Delta E$ (eV) | 4.00 | 4.00 | 3.69 | 3.85 | 4.23 | 4.92 |
| $\lambda$ (nm) | 310 | 310 | 336 | 322 | 293 | 252 |
| Oscillator strength | 0.31 | 0.32 | 0.01 | 0.62 | 0.56 | 0.16 |
| $\sigma(2PA)$ (GM) | 0.02 | 0.02 | 0.29 | 24.40 | 1.66 | 20.50 |
| D(3PA) | 1.28E+06 | 1.34E+06 | 5.96E+05 | 3.40E+07 | 7.38E+06 | 5.79E+06 |

Compared to CTL and DHE, the n-π* state of **5** possesses a modest two- and three-photon absorption. The π-π* state has a much larger two-photon cross-section (around 25 GM) and is also predicted to have relatively significant three-photon absorption properties, with a transition moment that is roughly 12 times larger than DHE/CTL. However, the close energetic proximity of the two states could potentially limit the fluorescence quantum yield of the molecule to some extent.

As shown in Fig. 1, the calculated natural transition orbitals (NTOs) of the n-π* state and the π-π* state of **5** are confined to the steroid ring system. The cholesterol analog **6** has two lowlying π-π* states but with a larger energetic separation (around 0.7 eV) compared to **5**. The absorption energy of the lowest state of **6** is predicted to be slightly blue-shifted compared to DHE and CTL (about 0.2 eV; Table 1). Both states are bright in a one-photon process, but the lower state possesses a higher oscillator strength. For two-photon absorption, the higher-lying state is predicted to be the more active state (at around 20 GM). The two states are both predicted to be active in a three-photon absorption process, with transition moments that are roughly five times larger than DHE/CTL. These results are important predictors of the optical properties of **5** and **6** for their future application in live-cell imaging of sterol transport. Particularly the much higher two- and three-photon absorption cross-section of **5** and **6** compared to DHE or CTL is promising, since multiphoton imaging of the latter sterols was found to give new insight into sterol distribution in model and cell membranes but was plagued by low signal-to-noise ratios.^[9b, 13]^

**Figure 1.**
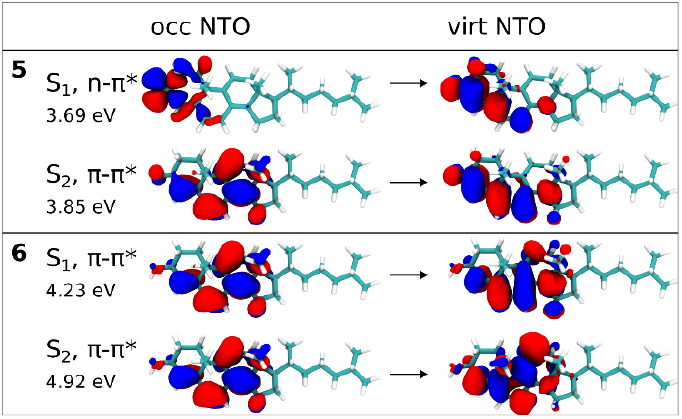
Electronic transitions energies and natural transition orbitals (NTOs) for **5** and **6**.

### Fluorescence properties of 5 and 6 in solvents

The fluorescence properties of **5** and **6** were studied in solvents of different polarity. The spectra reflect an approximately 10-fold higher fluorescence intensity of **5** than **6** (Fig. 2). Supporting this notion, we found a quantum yield of 25.6% for **5** but only 1.8% for **6**, both measured in ethanol at the respective absorption peak (see Fig. SI 1 and Table SI 1 for details). This difference may be due to the additional double bond of the 3’-carbonyl group in **5**, which makes **5** structurally more like a pentaene compared to **6**. The emission of both probes strongly depends on solvent properties; as there is a bathochromic shift between the solvents methanol, ethanol and isopropanol (Fig. 2, see also Fig. SI 2 showing the spectra normalized to the fluorescence maxima). This pronounced solvatochromism of both molecules is likely a consequence of their large change in dipole moment upon excitation. We did not consider in these experiments the spectra in chloroform owing to a sensitivity of the probes (especially for **6**) towards acids. Chloroform can easily form HCl which protonates the alcohol at the C3 position of **6**. Due to the double bonds of the molecule, this may cause a chemical reaction in that the alcohol is eliminated as H_2_O. Therefore, we note that these molecules should not be dissolved in chloroform which is often used as solvent for cholesterol.

**Figure 2.**
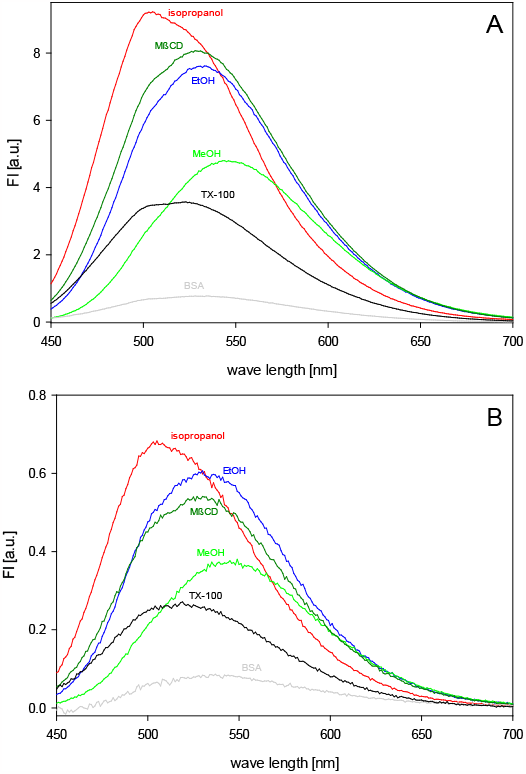
Fluorescence properties of **5** (A) and **6** (B) in different solvents. The fluorescent sterols were given from an ethanolic stock solution to the respective solvents and solutions (final concentration 10 μM): isopropanol (red line), ethanol (blue line), methanol (green line), Triton X-100 (0.7%(v/v)) (black line), methyl-β-cyclodextrin (1 mM) (dark green line), and bovine serum albumin (1%v/v)) (gray line). Fluorescence spectra (λ_ex_ = 396 nm) were recorded at room temperature. From the spectra, the spectra of the respective solvents/solutions were subtracted.

Additionally, fluorescence properties of the analogs were measured in the presence of Triton X-100, methyl-ß-cyclodextrin (MßCD) and bovine serum albumin (BSA) resulting in differently pronounced fluorescence spectra (Fig. 2 and Fig. SI 2). Triton X-100 is a detergent forming micelles in aqueous buffers into which the analogs are incorporated. Water-soluble cyclodextrins (CDs), especially methyl-β-cyclodextrin (MβCD), are able to form complexes with endogenous cholesterol, which has been used e.g. to manipulate the level of cholesterol and cholesterol analogs in model and biological membranes.^[25]^ The significant intensity in the spectra measured for **5** and **6** in MßCD show that both analogs also effectively bind to this CD which offers the possibility to label biological membranes with these analogs.

Albumins interact with a wide variety of hydrophobic ligands and serve physiologically in the serum as transporter for a variety of compounds e.g. fatty acids. It has been shown that BSA also binds cholesterol^[26]^ which underlines its role in cholesterol efflux and transfer between cells and lipoproteins.^[27]^ The weak fluorescence spectra measured for **5** and **6** in the presence of BSA, indicate that albumin also shows some affinity for the analogs. Comparing the position of the fluorescence maxima measured in the different solutions with those of the solvents allows to draw conclusions about the respective polarity around the analogs (see Fig. SI 2). From that, it can be postulated that **5** and **6** solubilized in Triton X-100 sense a polarity which is between that of isopropanol and ethanol, whereas when bound to MßCD or BSA, the polarity of their immediate environment is similar to that of ethanol.

Given that cholesterol has a very low water solubility and forms readily aggregates in aqueous solution^[28]^, we also studied the self-aggregation behavior of both sterol probes. We found in mixtures of water and ethanol that the fluorescence of **5** and **6** is strongly quenched, the higher the volume fraction of water is (Fig. SI 3). Additionally, in the emission of **6** a new band with a peak at 442 nm appeared at 75 % v/v water and above. The excitation spectra of both probes additionally revealed a shoulder peak between 310-350 nm, whose relative contribution to the overall intensity was higher in water than in ethanol. Using electronic structure calculations of dimers of either **5** or **6**, we found evidence for strong exciton splitting at short distances for both sterol probes with energy gaps of 0.062 and 0.053 eV, respectively (Fig. SI 4). Here, the resulting S1 state is dark (i.e., with close to zero oscillator strength), but in contrast, the resulting S2 state is bright with about twice the intensity of the isolated gas-phase state. Additional calculations including the polarized continuum model (to simulate environmental effects) revealed that the character of the states stays the same, and there is only about a 0.4 nm shift in the lowest excited state between water/ethanol (not shown). Thus, the experimentally observed differences in fluorescence in water solvent compared to ethanol solvent are not due to the difference in dielectric constants between either solvents. MD simulations additionally revealed that both **5** and **6** readily forms aggregates in water and in water/ethanol mixtures but not in pure ethanol (Fig. SI 5). Together, these results support the experimental observations, namely that both sterol probes self-aggregate in aqueous solution which causes a strong reduction in their fluorescence compared to less polar environments.

### Effect of 5 and 6 on the fatty acyl chains of phospholipids

One of the key properties of cholesterol is its ability to order acyl chains and thereby condense phospholipid membranes in the fluid phase. To determine to which extent **5** and **6** can mimic this effect of cholesterol, we carried out MD simulations of 1-palmitoyl-2-oleoyl-*sn*-glycero-3-phosphocholine (POPC) membranes containing either cholesterol, **5** or **6** (Fig. 3A, B). We found that both sterol probes increase the acyl chain order of both, the *sn-1* and *sn-2* chain of POPC, albeit to a lower degree than cholesterol. For both acyl chains, **6** was much more efficient than **5** in ordering along the entire carbon chains. To follow up on these observations, ^2^H NMR measurements of deuterated POPC-*d*_31_ were performed from which the smoothed order parameter of the *sn-1* chain was calculated (Fig. 3C). These experiments confirm the observations from MD simulations, namely that **6** can increase the acyl chain ordering of POPC, but not to the same extent as cholesterol does. There was no impact of **5** on acyl chain mobility in these ^2^H NMR experiments, suggesting that its impact on chain ordering is neglectable on the NMR time scale. We conclude from these studies, that the 3’-hydroxy group is an essential property for mediating cholesterol-like properties of a fluorescent cholesterol analog. The conjugated double bond system reduces the ability to pack membrane lipids, but to a smaller degree.

**Figure 3.**
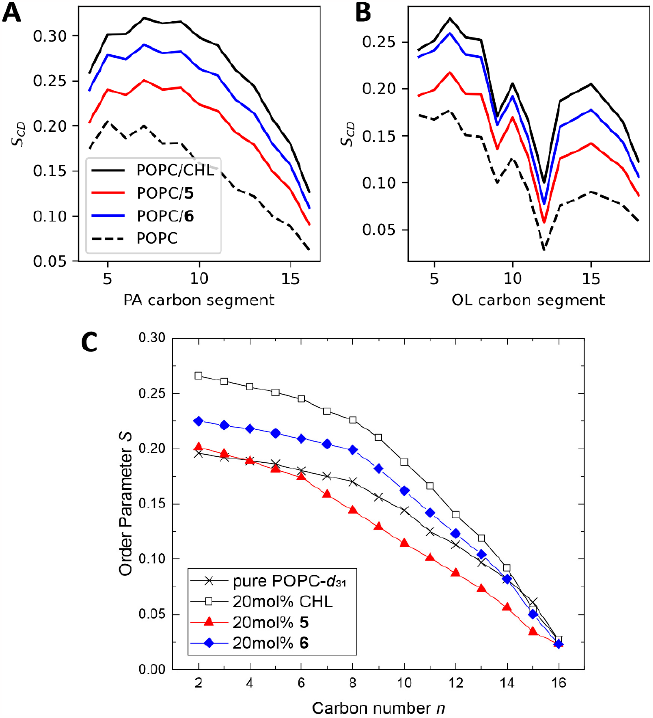
Influence of **5** and **6** on the phospholipid acyl chain ordering of POPC compared to that of cholesterol. Smoothed order parameters of the *sn-1* (A, C) or *sn-2* (B) chain are shown calculated from MD simulations (A, B) or obtained from ^2^H NMR measurements (C) without and with 20 mol% cholesterol, **5** or **6**.

### MD simulations of the membrane properties of 5 and 6

To further explore the structural requirements for membrane organization of the novel sterol analogs, the electron density profile and hydrogen-bonding capacity to interfacial water was analyzed from MD simulations (Fig. 4A). We found that membranes containing **6** but not **5** have a similar thickness compared to those with cholesterol. This supports the conclusion, that **6** can condense the lipid bilayer similar to cholesterol, which is known to increase the bilayer thickness at the expense of a reduced membrane area.^[29]^ In contrast, the 3’-carbonyl group of **5** causes reduced membrane packing, which is in line with previous observations on 3’-carbonyl cholesterol derivatives.^[30]^ Also, the capacity of forming hydrogen bonds to water is only slightly lower for **6** compared to cholesterol but much reduced for **5** (Fig. 4B). This capacity of hydrogen bonding is important for maintaining the structural integrity of membranes, partially for offsetting the large electrostatic repulsion between head groups and thereby contributing to the condensing effect of sterols.^[31]^ The fact that **6** has a comparable bonding capacity as cholesterol supports its close resemblance to cholesterol.

**Figure 4.**
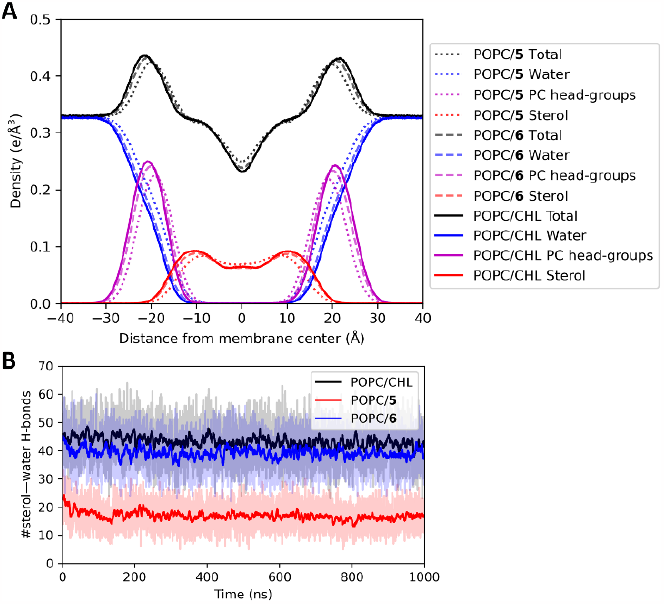
Electron density profiles and hydrogen-bonding of the sterol probes in POPC membranes. The figure shows electron density profiles (A) and hydrogen bonding to water molecules at the bilayer interface (B) for cholesterol (black line), **5** (red line) and **6** (blue line).

The ability of cholesterol and other sterols to condense and order lipid membranes is closely associated with the bilayer orientation of the sterol itself, with a low tilt angle being characteristic for a high capacity for membrane lipid packing. We calculated this tilt for the sterol probes and found that **5** but not **6** has a similarly confined and upright orientation in the membrane as cholesterol (Fig. 5). Together, these results clearly show, that the hydroxy, but not the keto form of the new sterol analogs, can mimic cholesterol’s membrane properties closely.

**Figure 5.**
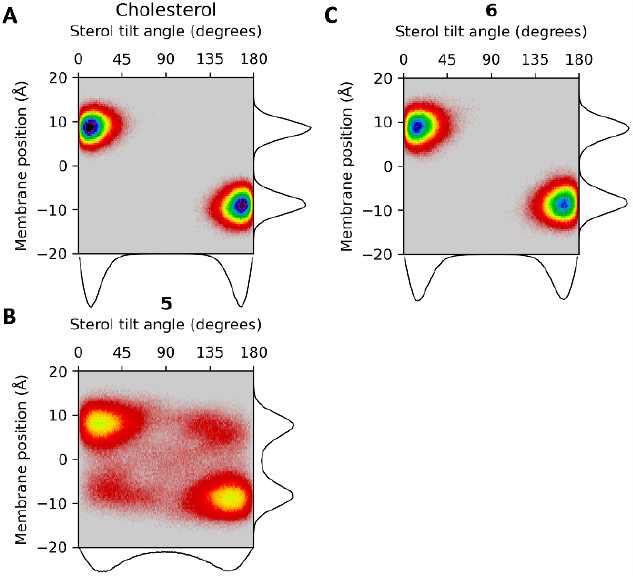
Sterol tilt angle distribution from MD simulations for cholesterol (A), **5** (B) and **6** (C).

### Fluorescence properties of 5 and 6 in membranes

Fluorescence excitation and emission spectra of the probes were recorded in multilamellar lipid vesicles (MLV) composed of POPC and POPC/cholesterol (2:1) and are shown in Fig. 6. They reveal that the fluorescence intensity of **6** amounts only to one tenth of that of **5** in each system. These results agree with the measurements in solvents (compare Fig. 2).

**Figure 6.**
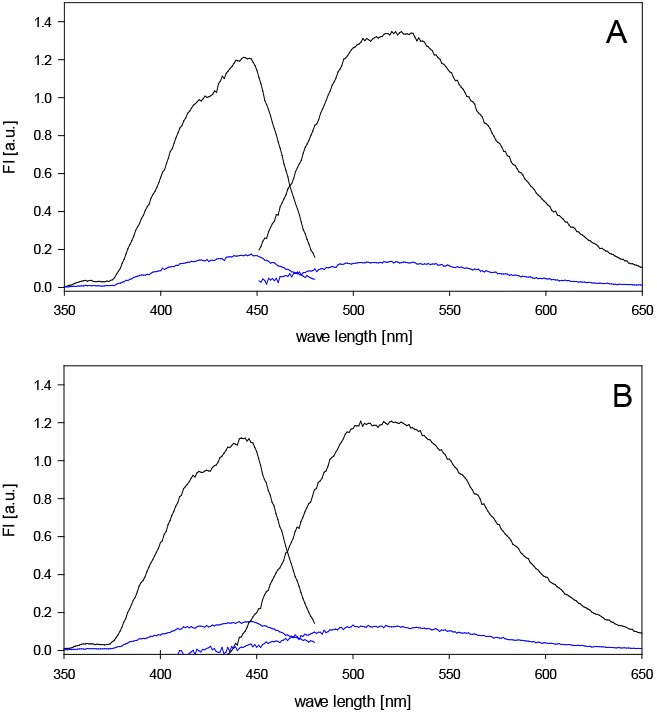
Fluorescence spectra of **5** and **6** in lipid membranes. Multilamellar vesicles (lipid concentration 200 μM) consisting of POPC (A) or POPC/cholesterol (2:1) (B) and either 10 μM **5** (black lines) or 10 μM **6** (blue lines) were prepared. The fluorescence excitation spectra (λ_em_ = 500 nm) and emission spectra (λ_ex_ = 396 nm) of the vesicles were recorded at room temperature. The spectra were corrected for scattering by subtracting the spectra of respective unlabelled lipid vesicles.

The maxima of the excitation and emission spectra of both probes are located at 443 nm (with a shoulder at 420 nm) and at 520 nm (with a shoulder appearing at 505 nm), respectively. Upon normalization of the spectra to their respective excitation and emission maxima, one sees that the spectra of both probes in membranes are very similar (Fig. SI 6).

The fluorescence lifetime of **5** and **6** in lipid membranes was determined in POPC MLV using a fluorescence lifetime spectrometer. The decay curves (Fig. SI 7) were fitted to a biexponential function giving time constants of 0.35 ns / 3.62 nm (**5**) and 0.27 ns / 3.59 ns (**6**). The component with the larger lifetime, which is very similar for both molecules, probably reflects analogs which are embedded as monomers in the membrane. The component with the much shorter lifetime could reflect a different population of the analogs in the membrane, for example sterol dimers and/or multimers experiencing fluorescence self-quenching. It has been shown that even cholesterol forms dimers and tetramers in lipid bilayers.^[32]^ From the above-mentioned time constants, an intensity weighted average fluorescence lifetime (τ_Av_) was calculated giving 2.77 ns +/- 0.07 ns for **5** and 2.28 +/- 0.05 ns for **6** (mean +/- SD, 3 measurements), revealing that both probes experience similar local environments in the lipid bilayer.

### Partitioning of 5 and 6 between various lipid phases

Cholesterol’s ability to condense membranes in the fluid phase together with its ‘fluidizing effect’ on gel-phase membranes can lead to macroscopic liquid-liquid phase separation, which is often studied in GUVs.^[33]^ Given the red-shifted emission of **5** and **6** compared to that of the established intrinsically fluorescent cholesterol and ergosterol analogs, CTL and DHE, respectively, we can image both, **5** and **6**, with conventional DAPI filters on a standard wide field microscope (Fig. 7). In ternary lipid mixtures consisting of POPC/N-stearoyl-*d-erythro*-sphingosyl-phosphorylcholine (SSM)/cholesterol and 5 mol% of either **5** or **6**, we found a significant partitioning of both sterol probes into the cholesterol-rich liquid-ordered (lo) phase compared to the liquid disordered (ld) phase labeled with the red-emitting marker 1,1′-dioctadecyl-3,3,3′,3′-tetramethylindo-carbocyanine perchlorate (DiIC18) (Fig. 7).^[33b]^ Quantification of sterol probe partitioning between both phases shows that **6** locates with higher preference to the lo phase (i.e. 62.7 +/- 9.8%), compared to **5**, which is on average distributed almost equally between the two phases, i.e. 55.3 +/- 7.1% are located in the lo phase. These results coincide very well with the differences between both cholesterol analogs with respect to their acyl chain ordering capacity (Fig. 3), hydrogen bonding capacity (Fig. 4B) and tilt in the bilayer (Fig. 5). The data underline, that the 3’-hydroxy group is essential for ‘good’ cholesterol-mimicking propensities of **6**. In addition, the flexibility of the ring system might be important, as we found by measuring the pseudo-valence angle between atoms 3, 9 and 16 of the steroids ring system slightly larger values for the mean and variance of this angle in **5** and **6** compared to cholesterol (Fig. SI 8).

**Figure 7.**
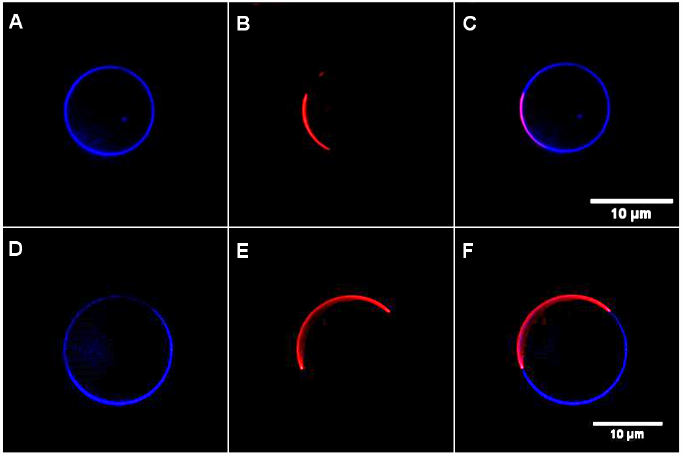
Fluorescence microscopic observation of domain partitioning of **5** and **6** in GUVs made of POPC/SSM/cholesterol, 0.1% of the ld marker DiIC18 and 5 mol% of **5** (A, B, C) or **6** (D, E, F). The analogs were imaged in the blue (DAPI) channel (A, D), while DiIC18 (B, E) was imaged in the red (Rhodamine) channel of a wide field microscope. Images were deconvolved before visualization. The color overlay is shown in C and F.

POPC GUVs were also used to measure the fluorescence lifetime of the analogs in lipid membranes using fluorescence lifetime imaging on a confocal microscope system FLIM (Fig. SI 9). Due to the lower quantum yield, the number of counts measured for **6** was too low preventing a determination of its lifetime. In contrast, for **5** enough counts could be measured allowing the calculation of reliable data. The decay curves were fitted to a biexponential function giving time constants of 4.07 ns +/- 0.22 ns and 1.15 ns +/- 0.06 ns. From these values, an intensity weighted average fluorescence lifetime (τ_Av_) of 3.12 ns +/- 0.07 ns (mean +/- SD, 4 measurements) was calculated, which is similar to the value obtained from bulk lifetime measurements on the spectrophotometer. Together, these results underline the potential of the novel sterol probes for advanced bioimaging applications.

## Conclusion

We present synthesis and characterization of two new fluorescent sterol analogs, one resembling endogenous cholesterol, the other 3 keto-sterol. Both molecules are distinguished by four conjugated double bonds in the sterol ring system which provides them with particular photo-physical properties. The improved photostability of both molecules allows their convenient imaging on conventional microscope systems which is a considerable advantage over other samples used so far. We show that the cholesterol-mimicking probe has comparable physicochemical and membrane properties as its endogenous counterpart. These probes open for many future applications in live-cell microscopy to investigate the trafficking of cholesterol in cells under normal and disease-related conditions.

## Supporting Information

The Supporting Information contains experimental procedures and supporting figures and tables. The authors have cited additional references within the Supporting Information.^[34-58]^

### Acknowledgements

This research was funded by the Deutsche Forschungsgemeinschaft (DFG), MU 1017/12-1 (P.M. and D.W. as Mercator Fellow), and WE1850-12/1 (P.W.) as well as by the Lundbeck Foundation (R366-2021-226) to DW. DW and JK acknowledge financial support from The Independent Research Fund Denmark – Natural Sciences (Grant ID DFF-7014-00050B).

## Entry for the Table of Contents

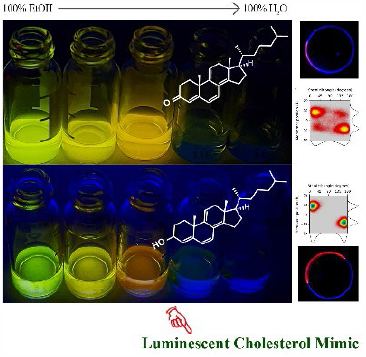

Novel intrinsically fluorescent sterol probes are synthesized and characterized with improved photophysical properties and close resemblance of the membrane properties of cholesterol. The analogs can be visualized using conventional microscope systems making them very useful for tracking cellular cholesterol trafficking.

