## Supplemental figures for "Synthesis and characterization of novel intrinsically fluorescent analogs of cholesterol with improved photophysical properties"

Supporting Information
©Wiley-VCH 2021
69451 Weinheim, Germany

### Synthesis and characterization of novel intrinsically fluorescent analogs of cholesterol with improved photophysical properties

*M. Lehmann, P. Reinholdt, M. Bashawat, H. A. Scheidt, S. Halder, D. di Prima, J. Kongsted*, P. Müller*, P. Wessig*, D. Wüstner**

**DOI: 10.1002/anie.2021XXXXX**

#### Table of Contents

Experimental Procedures page 2
Supporting Figures page 18
References page 27

#### Experimental Procedures

*Synthetic procedures and analytical data*

*General.* All reagents and solvents were procured from commercial suppliers and were utilized with additional purification if deemed necessary. Reactions were conducted under an inert atmosphere, either nitrogen or argon, when anhydrous solvents were employed, and they were monitored using thin-layer chromatography (TLC) on aluminum sheets coated with silica gel (SiO2-60, F254). Flash chromatography was executed on silica gel 60 (40–63 mm). Nuclear magnetic resonance (NMR) spectra were recorded using a Bruker Avance 300 spectrometer, with chemical shifts reported relative to solvent signals as follows: $^1$H NMR CDCl3: δ = 7.26 ppm, $CD_2Cl_2$: δ = 5.32 ppm and $^{13}$C NMR $CDCl_3$: δ = 77.0 ppm, $CDCl_2$: δ = 54.0 ppm. High-resolution mass spectrometry (HRMS) data were acquired using a quadrupole time-of-flight (TOF) mass spectrometer. Melting points were determined using an Elektrothermal 9100 melting point instrument. The IR spectra were recorded using the Spectrum Two IR spectrometer from PERKIN ELMER (UATR Two). The PERKIN ELMER SPECTRUM software was used. The vibration bands ($\tilde{\nu}$) were given without assignment in $cm^{-1}$.

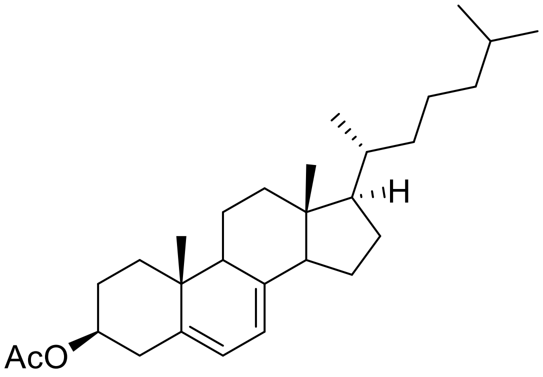


Compound **2**: (3S,10R,13R,17R)-10,13-Dimethyl-17-((R)-6-methylheptan-2-yl)-2,3,4,9,10,11,12,13,14,15,16,17-dodecahydro-1H-cyclopenta[a]phenanthren-3-yl acetate 2

3.00 g (7.08 mmol, 1.0 eq.) 7-dehydrocholesterol **1** was dissolved in 50 mL pyridine, cooled to 0°C and mixed with 1.01 mL (14.16 mmol, 2.0 eq.) acetyl chloride. The mixture was stirred overnight and heated to RT. After complete conversion, the reaction mixture was placed on 200 g ice, the precipitated white solid was dissolved with 100 mL DCM and the phases were separated. The organic phase was washed with 28% HCl, the phases were separated again and the organic phase was neutralised with $NaHCO_3$. The solution was finally dried over MgSO4, the solvent removed on the rotary evaporator and purified by FC (PE:EE, 20:1). 3.01 g (7.05 mmol) of the compound **2** could be prepared as a white solid with a yield of 100 %.
**$^1$H NMR:** (300 MHz, $CDCl_3$) δ 5.59 – 5.53 (m, 1H), 5.42 – 5.35 (m, 1H), 4.78 – 4.61 (m, 1H), 2.04 (s, 3H), 0.96 – 0.91 (m, 6H), 0.88 – 0.86 (m, 3H), 0.86 – 0.84 (m, 3H), 0.61 (s, 3H).
**$^{13}$C NMR**: (75 MHz, $CDCl_3$) δ 170.7, 141.7, 138.7, 120.4, 116.4, 73.0, 56.0, 54.6, 46.2, 43.1, 39.7, 39.3, 38.1, 37.2, 36.8, 36.3, 36.3, 28.3, 28.2, 24.0, 23.2, 23.0, 22.7, 21.6, 21.2, 19.0, 16.3, 12.0.
**IR:** (ATR, $cm^{-1}$) $\tilde{\nu}$ = 3430, 2951, 2931, 2870, 1735, 1467, 1365, 1242, 1031.
**Smp.:** 122 – 123 °C.
**HRMS**: [ESI] m/z calcd. for $C_{29}H_{46}O_2[M^+]$: 427.3576, found: 427.3560.

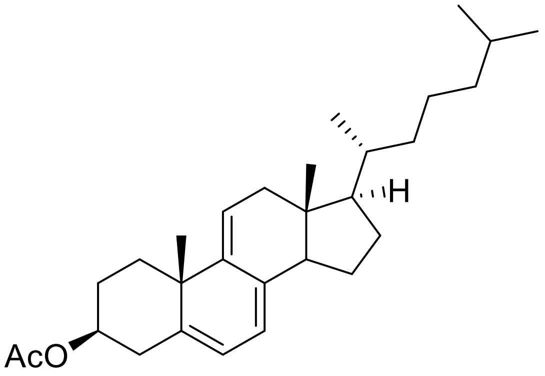

Compound **3**: (3S,10S,13R,17R)-10,13-Dimethyl-17-((R)-6-methylheptan-2-yl)-2,3,4,10,12,13,14,15,16,17-decahydro-1H-cyclopenta[a]phenanthren-3-yl acetate 3

2.27 g (5.32 mmol, 1.0 eq.) of the diene **2** were dissolved in 50 mL ethanol and 15 mL DCM. Then 5.09 g (19.97 mmol, 3.0 eq.) of mercuric acetate was added first, followed by 2.74 mL (48.0 mmol, 0.2 eq.) of acetic acid. The suspension was stirred for 48 h, the solid was filtered and the filtrate was washed several times with a 5% NaCl solution. The organic phase was dried over $MgSO_4$, the solvent was removed on a rotary evaporator and the crude product was purified by FC (PE:EE, 10:1). Compound **3** was obtained as a lightly yellow solid in a yield of 65% (1.48 g, 3.48 mmol).
**$^{1}$H NMR**: (300 MHz, $CDCl_3$) δ 5.71 – 5.64 (m, 1H), 5.54 – 5.45 (m, 1H), 5.44 – 5.35 (m, 1H), 4.71 – 4.60 (m, 1H), 2.03 (s, 3H), 1.25 (s, 3H), 0.92 (d, J = 6.3 Hz, 3H), 0.88 – 0.86 (m, 3H), 0.86 – 0.84 (m, 3H), 0.56 (s, 3H).
**$^{13}$C NMR:** (75 MHz, $CDCl_3$) δ 170.5, 144.1, 140.1, 135.8, 122.7, 119.2, 115.7, 74.2, 56.5, 51.1, 43.2, 42.4, 39.6, 39.5, 38.2, 37.6, 36.2, 36.1, 30.4, 28.6, 28.5, 28.2, 24.1, 23.0, 22.7, 21.5, 18.5, 11.5.
**IR**: (ATR, cm-1) $\tilde{\nu}$ == 2951.63, 2926.64, 2866.88, 2820.17, 1726.96, 1466.08, 1435.79, 1375.62, 1361.93, 1241.43.
**Smp.:** 84 – 85°C.
**HRMS**: [ESI] m/z calcd. for C22H40 $[M+H(-OAc)]^{\bullet+}$: 364.3152, found: 365.320.

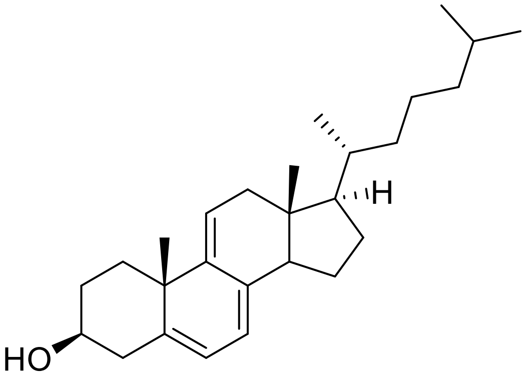

Compound **4**: (3S,10S,13R,17R)-10,13-Dimethyl-17-((R)-6-methylheptan-2-yl)-2,3,4,10,12,13,14,15,16,17-decahydro-1H-cyclopenta[a]phenanthren-3-ol 4

180.0 mg (423.9 µmol, 1.0 eq.) of the acetate **3** was dissolved in 50 mL MeOH, NaOH (5%) was added and the solution was heated for approx. 1 h under reflux. After completion of the reaction, 30 mL of water and 30 mL of DCM were added, the phases separated and the aqueous phase extracted again with 30 mL of DCM. The combined organic phases were then dried over $MgSO_4$ and the solvent removed. After column chromatographic purification (PE:EE, 20:1) **4** (121.0 mg, 316.2 µmol, 75%) was obtained as a lightly yellow solid.
**$^{1}$H NMR:** (300 MHz, $CDCl_3$) δ 5.71 – 5.64 (m, 1H), 5.54 – 5.48 (m, 1H), 5.43 – 5.36 (m, 1H), 3.66 – 3.54 (m, 1H), 1.24 (s, 2H), 0.92 (d, J = 6.3 Hz, 4H), 0.88 (d, J = 1.4 Hz, 3H), 0.85 (d, J = 1.4 Hz, 2H), 0.56 (s, 3H).
**$^{13}$C NMR:** (75 MHz, $CDCl_3$) δ 162.5, 144.3, 141.4, 135.62, 122.7, 118.4, 115.8, 72.4, 56.5, 51.1, 43.2, 42.4, 41.7, 39.6, 39.5, 38.5, 36.2, 36.1, 32.4, 30.6, 28.7, 28.2, 24.1, 23.0, 22.7, 18.6, 11.5.
**IR:** (ATR, cm-1) $\tilde{\nu}$ = 3365, 2954, 2932, 2868, 1464, 1431, 1376, 1358, 1048, 833.
**Smp.:** 104 – 105°C.
**HRMS:** [ESI] m/z calcd. for $C_{27}H_{41}$ [M+(-OH)]: 365.3208, found: 365.3191.

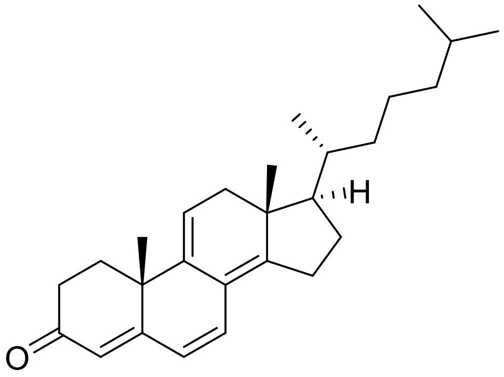

Compound **5**: (10S,13R,17R)-10,13-Dimethyl-17-((R)-6-methylheptan-2-yl)-1,2,10,12,13,15,16,17-octahydro-3H-cyclopenta[a]phenanthren-3-one 5

5 mL dry DCM was cooled to -78°C and 0.179 mL (2.09 mmol, 10.0 eq.) oxalyl chloride was added. Subsequently, 0.297 mL (4.18 mmol, 20.0 eq.) DMSO was added dropwise. The solution was stirred for 30 min and then 80 mg (209.1 µmol, 1.0 eq.) of compound **4** dissolved in 2 mL DCM was added. The solution was stirred for another hour at -78°C. Subsequently, 0.588 mL (4.18 mmol, 20.0 eq.) of triethylamine was slowly added. The reaction mixture was stirred for another hour while warming to room temperature. The reaction was then terminated by the addition of water (5 mL) and the phases separated. The aqueous phase was extracted with DCM (3x 10 mL). The combined organic phases were dried over magnesium sulphate and the solvent was removed on the rotary evaporator. The crude product was purified by flash chromatography (PE:EE, 10:1). Compound **5** was obtained as orange oil in a yield of 61.8 mg (163.2 µmol, 78%).

**$^{1}$H NMR:** (400 MHz, $CDCl_3$) δ 6.58 (d, J = 9.5 Hz, 1H), 6.07 (d, J = 9.5 Hz, 1H), 5.79 (s, 1H), 5.52 (dd, J = 6.8, 2.3 Hz, 1H), 1.27 (s, 3H), 0.95 (d, J = 6.3 Hz, 3H), 0.90 (s, 3H), 0.88 (d, J = 1.6 Hz, 3H), 0.87 (d, J = 1.6 Hz, 3H).

**$^{13}$C NMR:** (101 MHz, $CDCl_3$) δ 199.2, 162.5, 157.2, 138.7, 131.3, 124.1, 123.6, 122.2, 116.2, 57.3, 44.2, 39.6, 38.1, 37.7, 36.1, 34.3, 34.1, 32.3, 28.3, 28.2, 28.0, 26.6, 23.9, 23.0, 22.7, 18.9, 16.04, 0.1.

**IR:** (ATR, cm-1) $\tilde{\nu}$ = 2956, 2930, 2867, 1651, 1583, 1464, 1327, 1228, 873, 757.

**HRMS:** [EI] m/z calcd. for $C_{27}H_{38}O^{+}$[M+]: 378.2923, found: 378.2922.

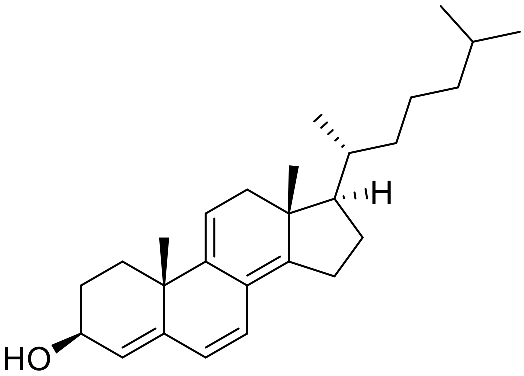

Compound **6**: (3S,10S,13R,17R)-10,13-Dimethyl-17-((R)-6-methylheptan-2-yl)-2,3,10,12,13,15,16,17-octahydro-1H-cyclopenta[a]phenanthren-3-ol 6

250.0 mg (660.3 µmol, 1.0 eq.) of the enone **5** and 270.6 mg (726.3 µmol, 1.1 eq.) cerium(III) chloride were dissolved in a mixture of THF:MeOH (20 mL, 3:1) and cooled to 0°C. The reaction mixture was mixed with 25.0 mg (660.3 µmol, 1.0 eq.) $NaBH_4$ and stirred for approx. 1 h. To the reaction mixture was added 25.0 mg (660.3 µmol, 1.0 eq.) $NaBH_4$ and stirred for about 1 h. The reaction was stopped, and the reaction was completed. After complete conversion, the reaction was terminated by adding 10 mL water, the phases were separated and the aqueous phase was extracted several times with DCM. The combined organic phases were dried over $MgSO_4$, the solvent removed and the crude product purified by HPLC (DCM:MeOH, 100:1). Compound **6** was obtained as a yellow solid in a yield of 56% (140.0 mg, 367.8 µmol).

**$^{1}$H NMR:** (400 MHz, $CD_2Cl_2$) δ 6.18 (d, J = 9.7 Hz, 1H), 5.91 (d, J = 9.7 Hz, 1H), 5.45 (d, J = 2.5 Hz, 1H), 5.36 (dd, J = 6.9, 2.2 Hz, 2H), 4.32 – 4.23 (m, 1H), 1.16 (s, 3H), 0.94 (d, J = 6.4 Hz, 3H), 0.88 (d, J = 1.3 Hz, 3H), 0.87 (s, 6H).

**$^{13}$C NMR:** (101 MHz, $CD_2Cl_2$) δ 151.7, 143.4, 141.5, 128.1, 125.4, 124.3, 122.8, 114.4, 68.6, 57.8, 44.2, 40.1, 38.7, 37.7, 36.6, 34.6, 32.5, 30.1, 29.5, 28.8, 28.6, 26.6, 24.3, 23.1, 22.9, 19.2, 16.3.

**IR:** (ATR, cm-1) $\tilde{\nu}$ = 3544, 2956, 2927, 2870, 2862, 1463, 1373, 1243, 1028, 864.

**Smp.:** 95°C.

**HRMS:** [EI] m/z calcd. for $C_{27}H_{40}O^{+}$[M+]: 380.3079, found: 380.3075.

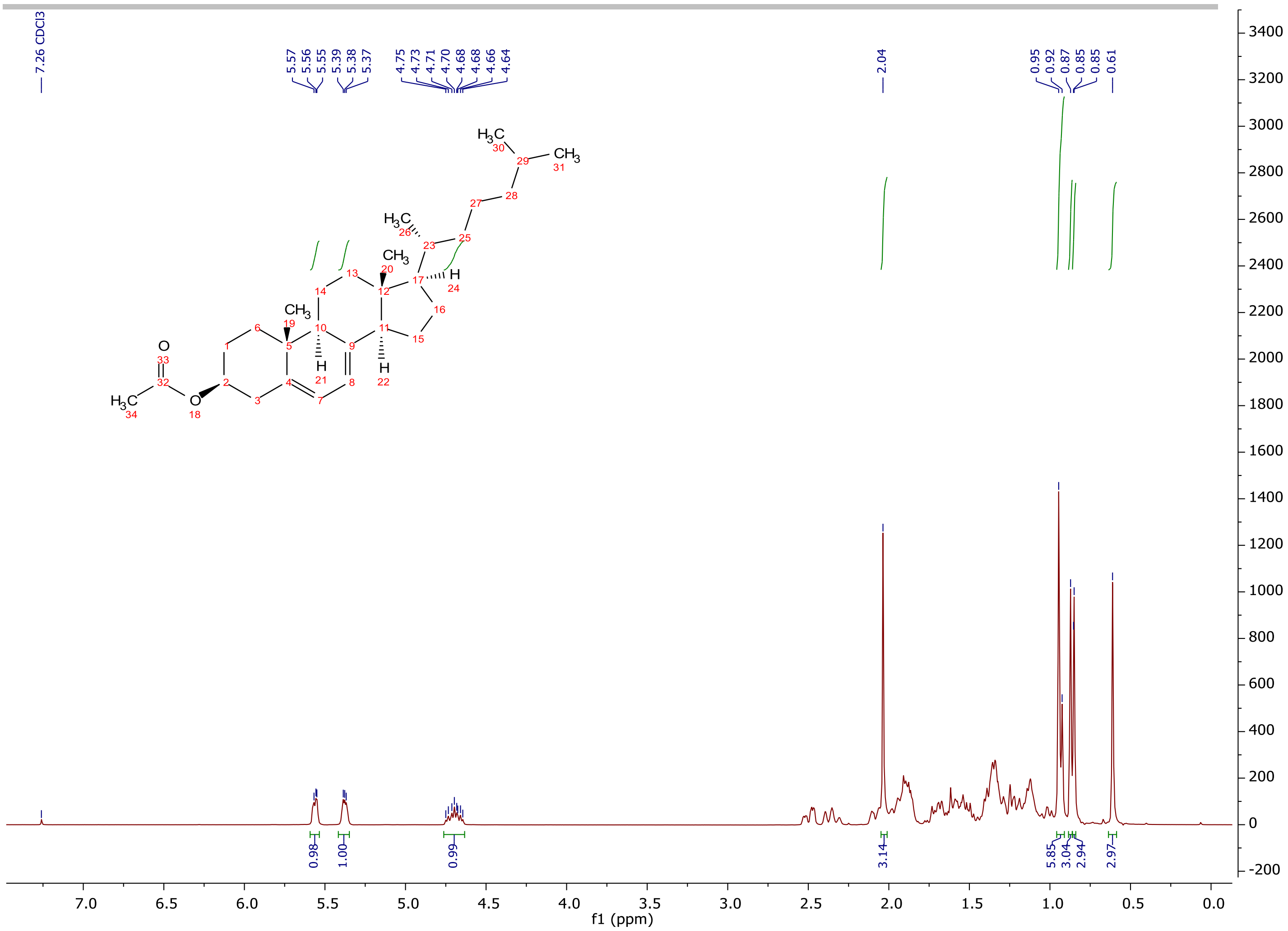
7.26 CDCl3
5.57
5.56
5.55
5.39
5.38
5.37
4.75
4.73
4.71
4.70
4.68
4.68
4.66
4.64
2.04
0.95
0.92
0.87
0.85
0.85
0.61
0.98
1.00
0.99
3.14
5.85
3.04
2.94
2.97
f1 (ppm)

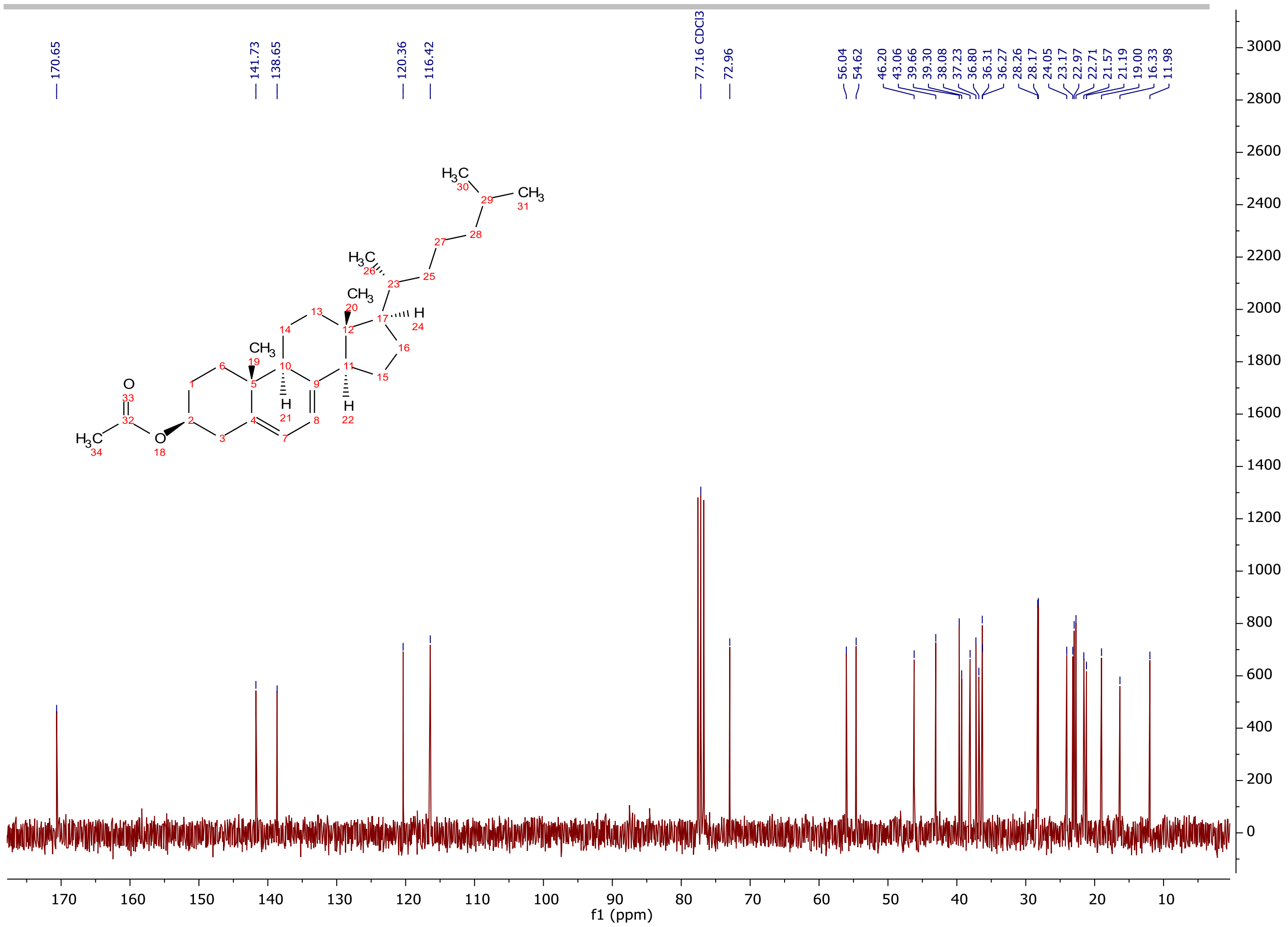
170.65
141.73
138.65
120.36
116.42
77.16 CDCl3
72.96
56.04
54.62
46.20
43.06
39.66
39.30
38.08
37.23
36.80
36.31
36.27
28.26
28.17
24.05
23.17
22.97
22.71
21.57
21.19
19.00
16.33
11.98
f1 (ppm)

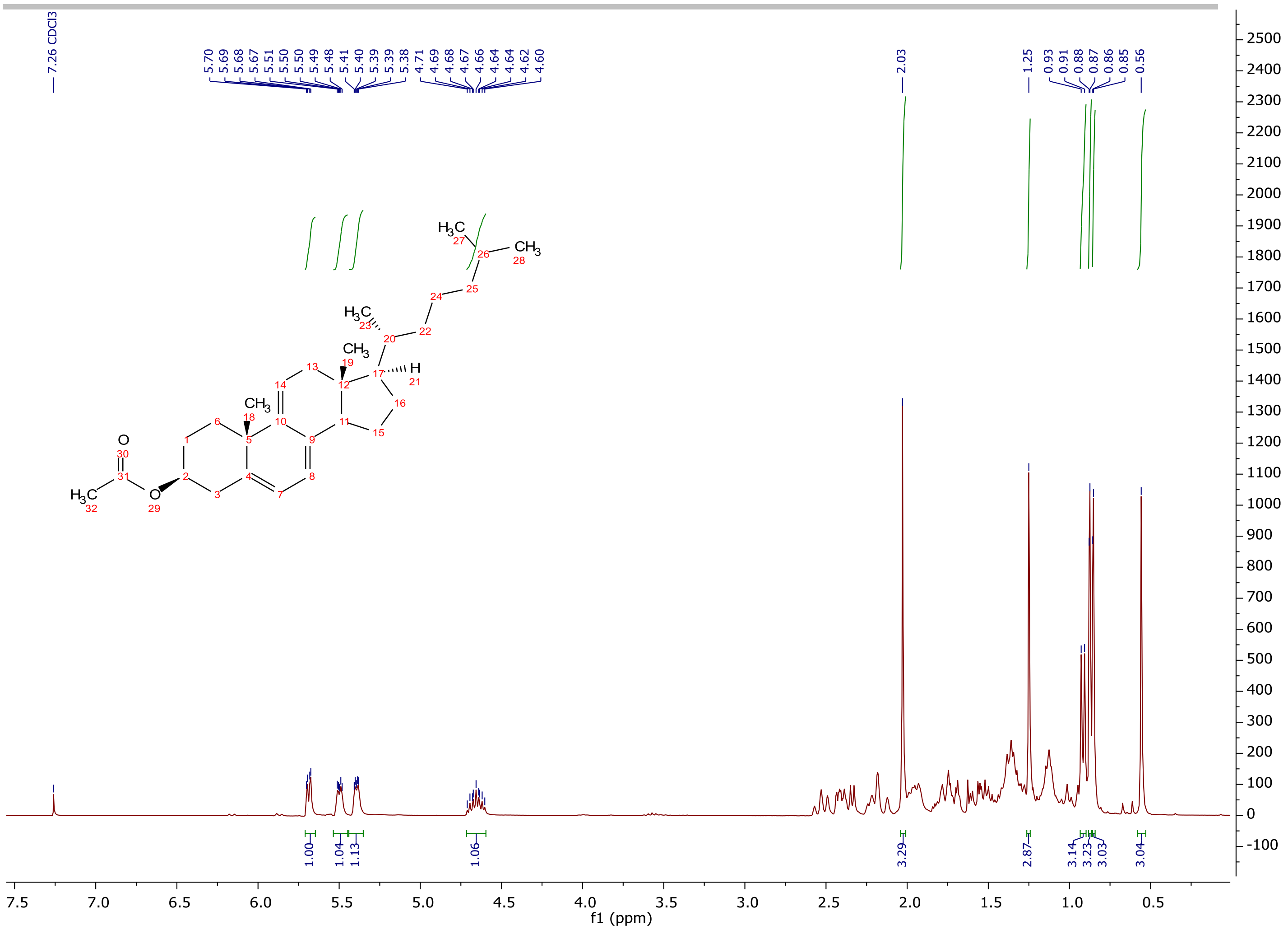

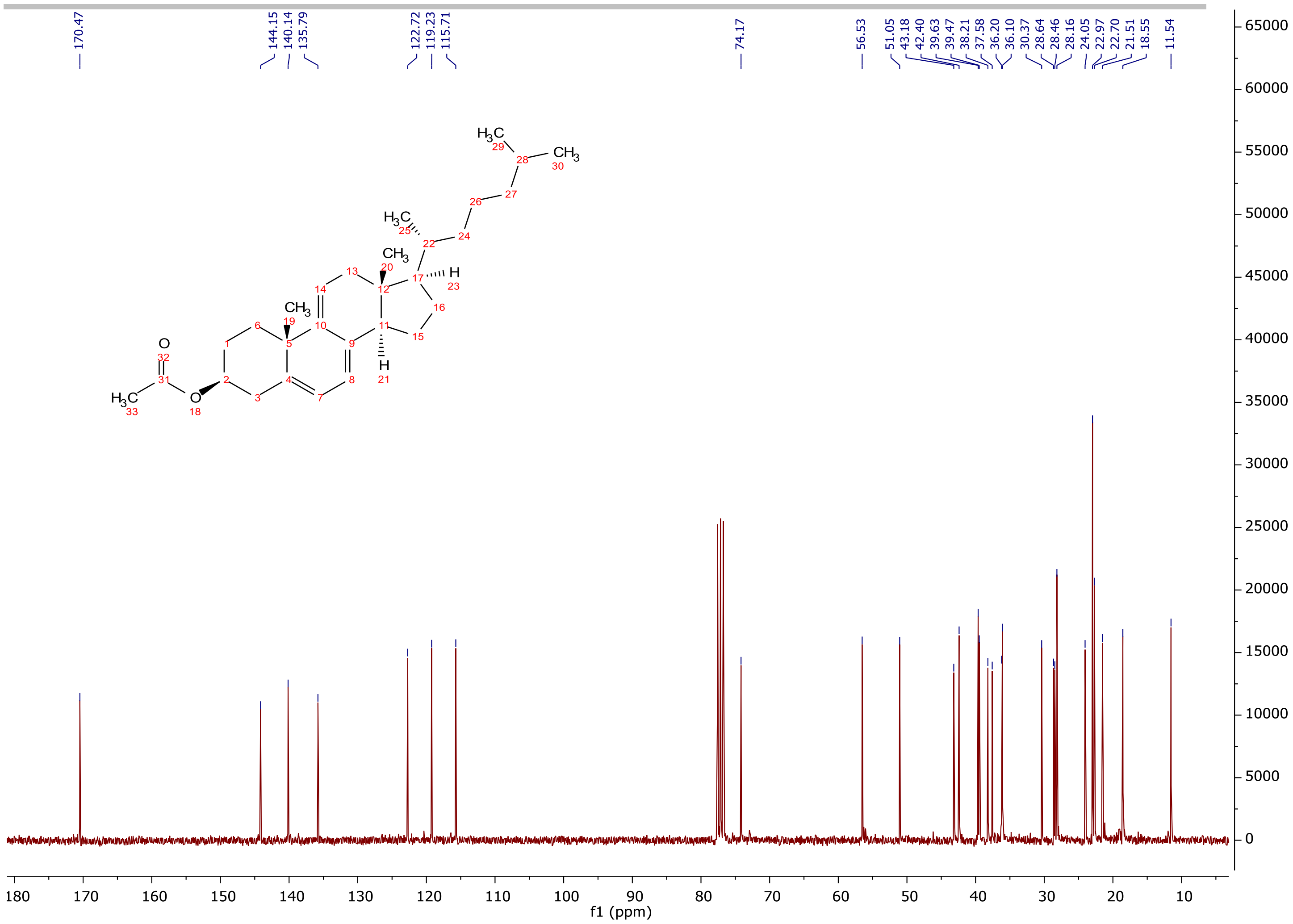
170.47
144.15
140.14
135.79
122.72
119.23
115.71
74.17
56.53
51.05
43.18
42.40
39.63
39.47
38.21
37.58
36.20
36.10
30.37
28.64
28.46
28.16
24.05
22.97
22.70
21.51
18.55
11.54
f1 (ppm)

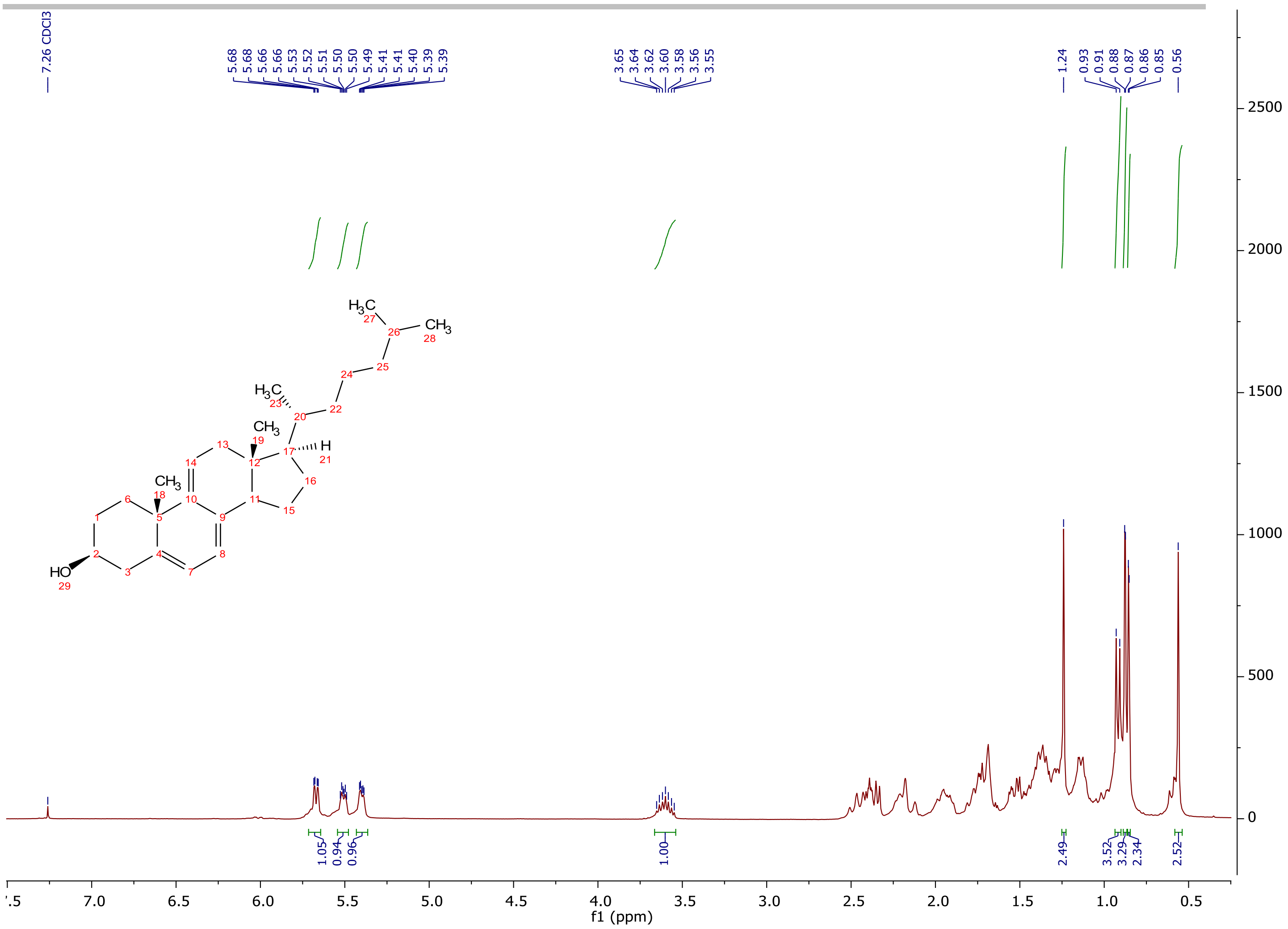
7.26 CDCl3
5.68
5.68
5.66
5.66
5.53
5.52
5.51
5.50
5.50
5.49
5.41
5.41
5.40
5.39
5.39
3.65
3.64
3.62
3.60
3.58
3.56
3.55
1.24
0.93
0.91
0.88
0.87
0.86
0.85
0.56
1.05
0.94
0.96
1.00
2.49
3.52
3.29
2.34
2.52
f1 (ppm)
HO
CH3
H3C
H
2500
2000
1500
1000
500
0

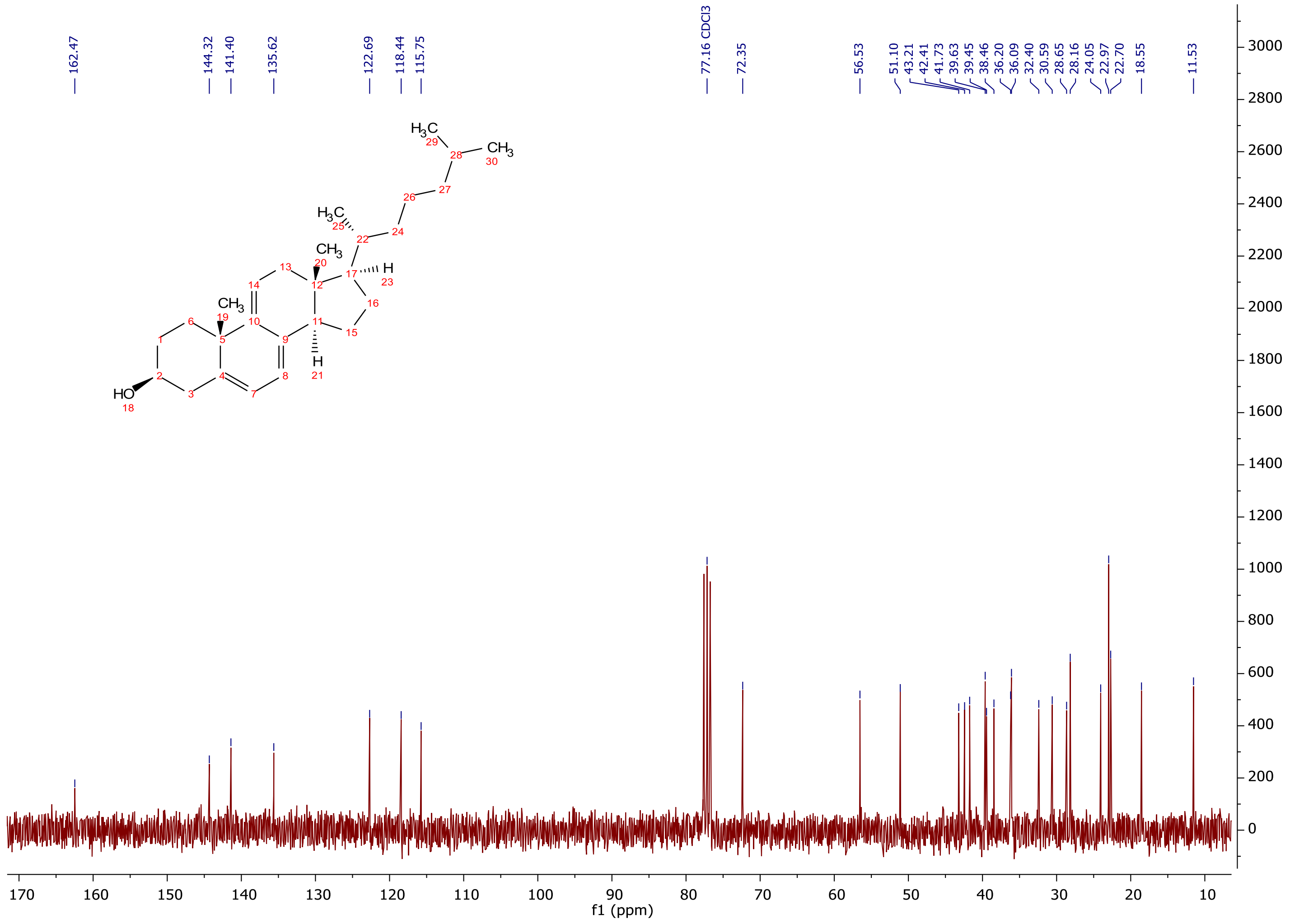
162.47
144.32
141.40
135.62
122.69
118.44
115.75
77.16 CDCl3
72.35
56.53
51.10
43.21
42.41
41.73
39.63
39.45
38.46
36.20
36.09
32.40
30.59
28.65
28.16
24.05
22.97
22.70
18.55
11.53
HO
CH3
H3C
H
f1 (ppm)

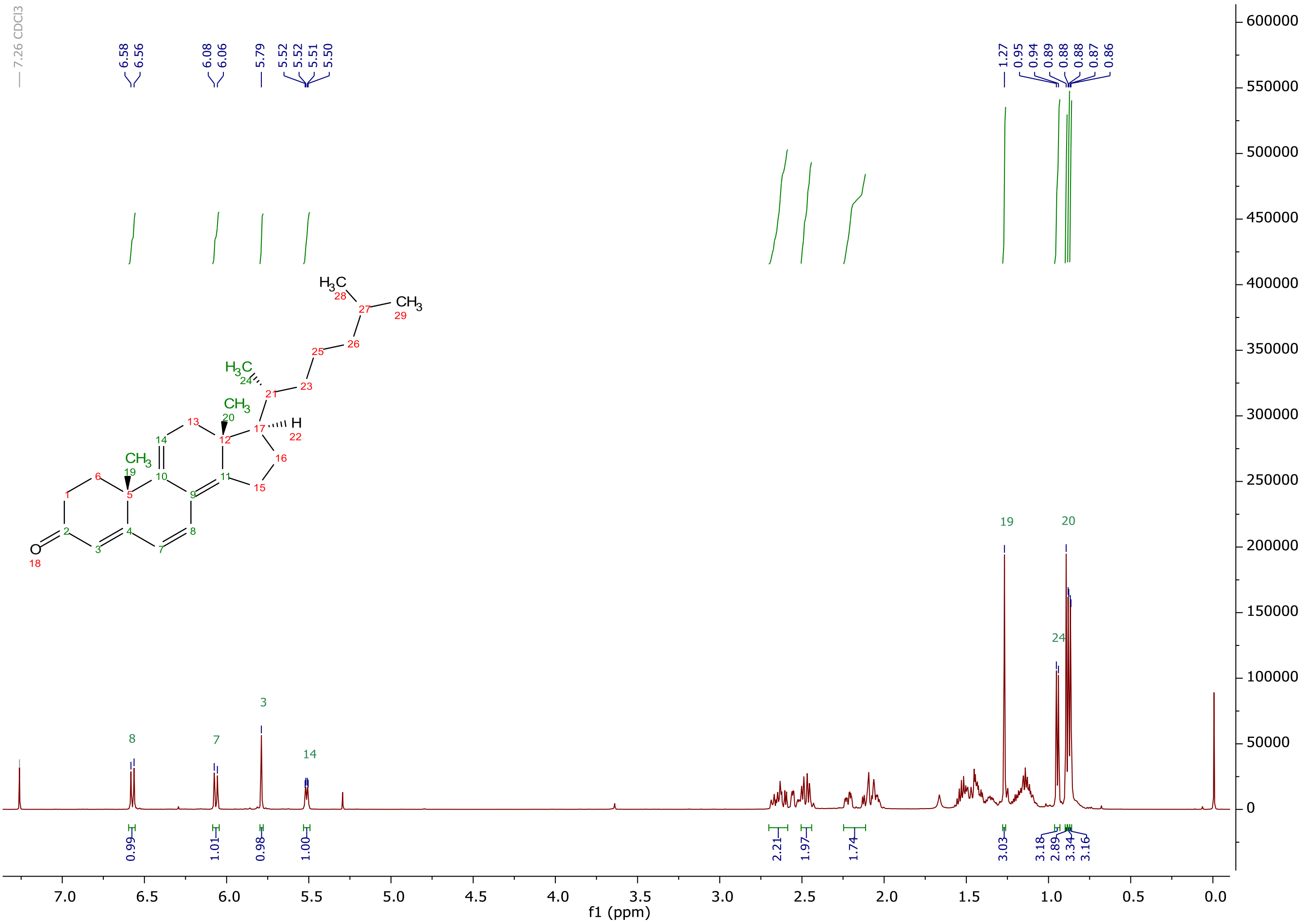
7.26 CDCl3
6.58
6.56
6.08
6.06
5.79
5.52
5.52
5.51
5.50
1.27
0.95
0.94
0.89
0.88
0.88
0.87
0.86
0.99
1.01
0.98
1.00
2.21
1.97
1.74
3.03
3.18
2.89
3.34
3.16
8
7
3
14
19
20
24
f1 (ppm)

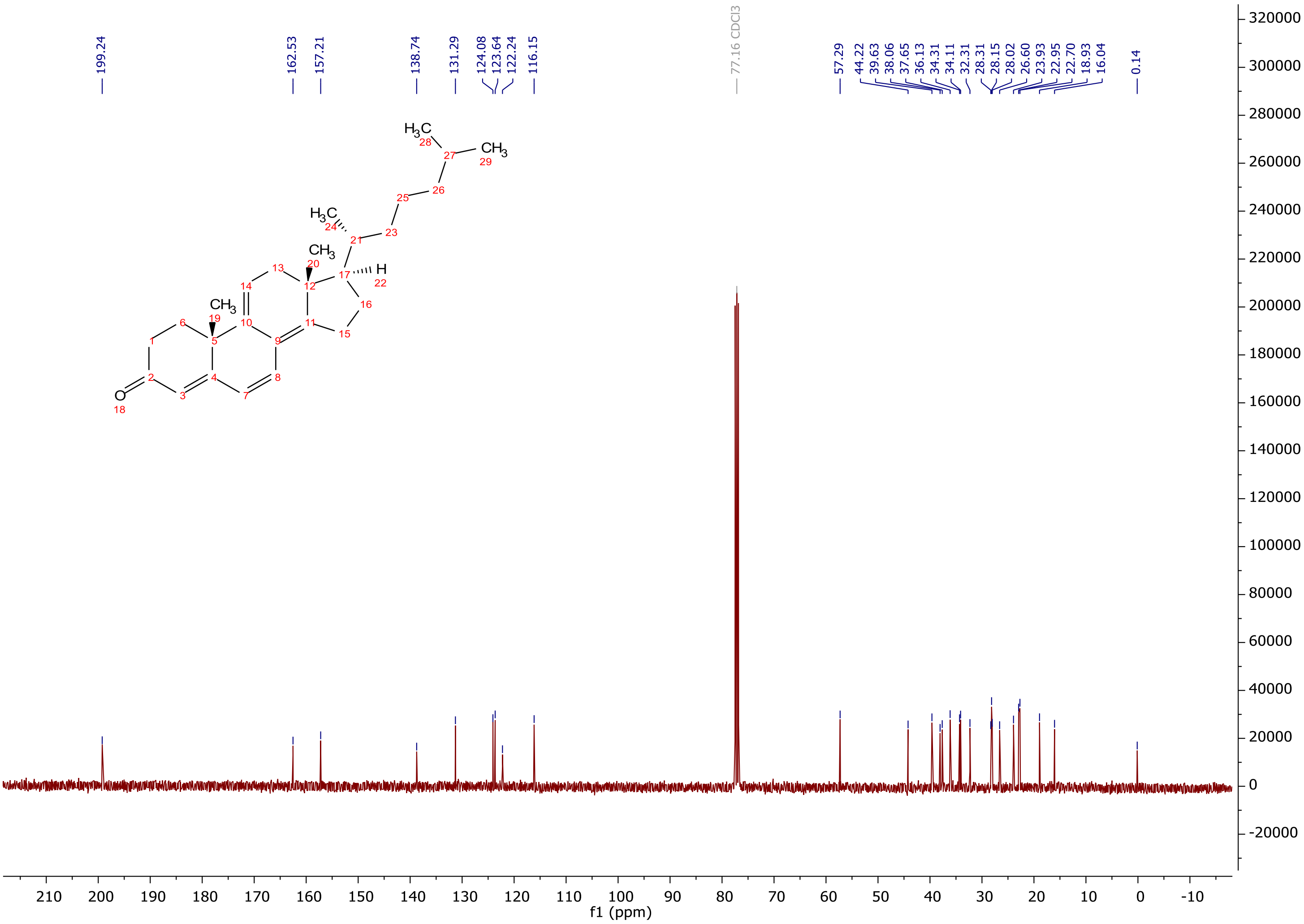
199.24
162.53
157.21
138.74
131.29
124.08
123.64
122.24
116.15
77.16 CDCl3
57.29
44.22
39.63
38.06
37.65
36.13
34.31
34.11
32.31
28.31
28.15
28.02
26.60
23.93
22.95
22.70
18.93
16.04
0.14
f1 (ppm)

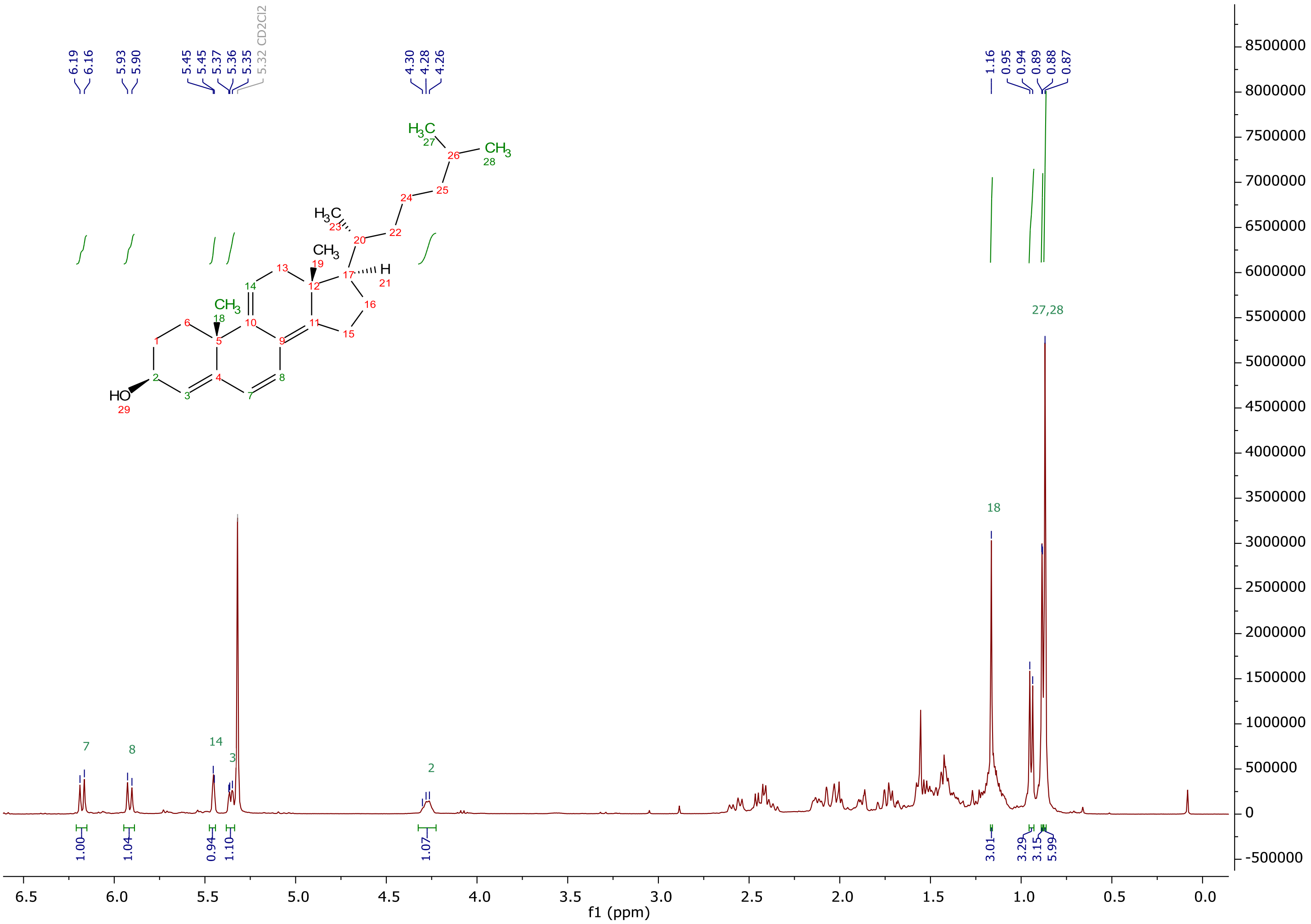
6.19
6.16
5.93
5.90
5.45
5.45
5.37
5.36
5.35
5.32 CD2Cl2
4.30
4.28
4.26
1.16
0.95
0.94
0.89
0.88
0.87
H3C
CH3
HO
27,28
18
7
8
14
3
2
1.00
1.04
0.94
1.10
1.07
3.01
3.29
3.15
5.99
f1 (ppm)

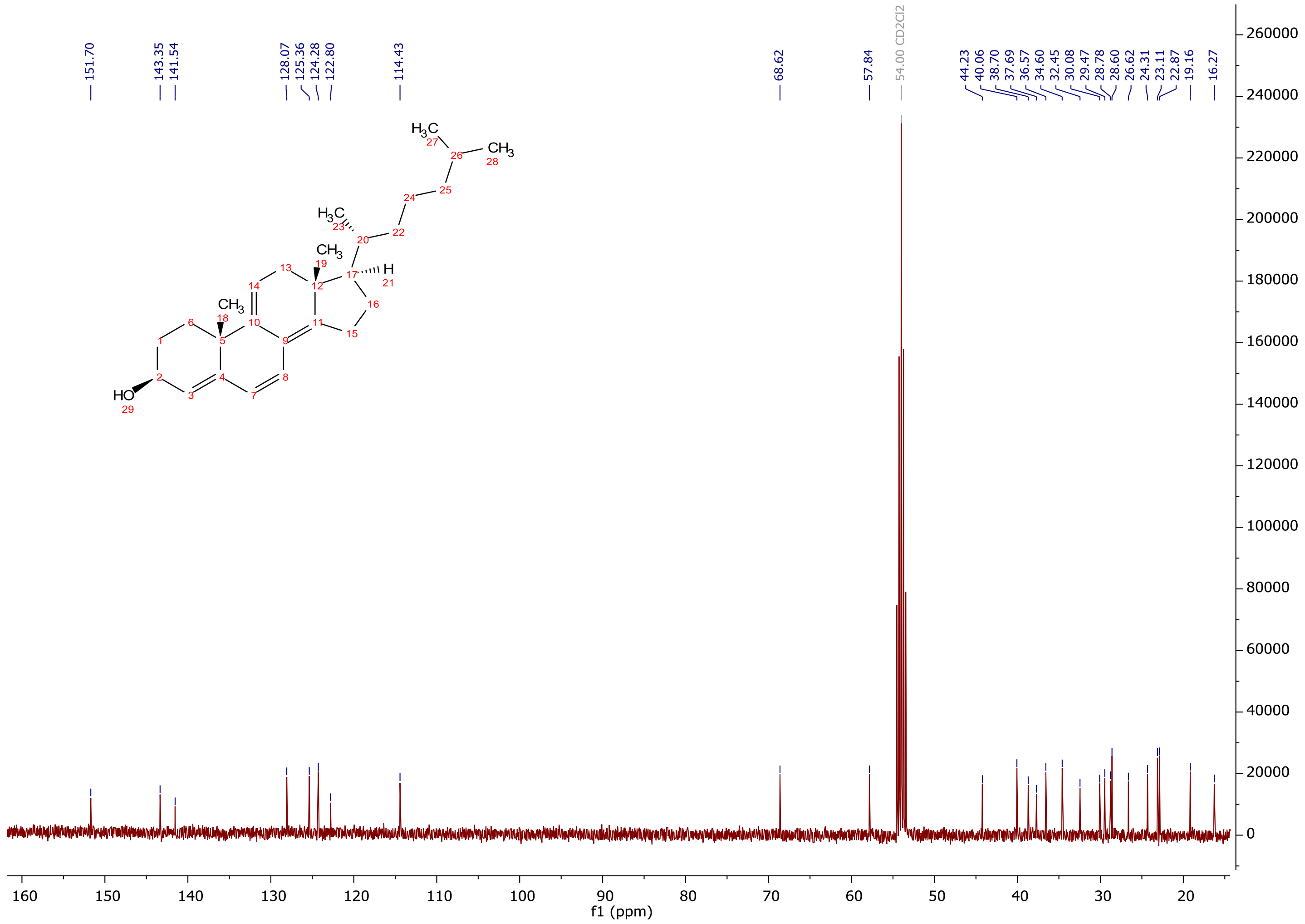
151.70
143.35
141.54
128.07
125.36
124.28
122.80
114.43
68.62
57.84
54.00 CD2Cl2
44.23
40.06
38.70
37.69
36.57
34.60
32.45
30.08
29.47
28.78
28.60
26.62
24.31
23.11
22.87
19.16
16.27
HO
CH3
H3C
H
f1 (ppm)

*Chemicals:* The fluorescent stain 1,1'-dioctadecyl-3,3,3',3'-tetramethylindocarbocyanine perchlorate (DiIC18) was purchased from Thermo Fisher and 1-palmitoyl-$d_{31}$-2-oleoyl-*sn*-glycero-3-phosphocholine (POPC-$d_{31}$) and cholesterol from Avanti Polar Lipids, Inc. (Alabaster, Al). All other chemicals were obtained from Sigma-Aldrich (Taufkirchen, Germany).

*Preparation of multilamellar vesicles (MLVs):* Aliquots of lipids dissolved in EtOH were transferred to a glass tube, dried under nitrogen and high vacuum and resuspended in a small volume of ethanol (to resolve the lipids from the glass tube) and, subsequently, in HEPES buffered salt solution (HBS, 10 mM HEPES and 145 mM NaCl, pH 7.4) (final ethanol concentration was below 1% (v/v)). By subsequent vortexing of the samples, MLVs were formed.[1] Vesicles were stored at 4°C and used within the same day.

*Preparation of giant unilamellar vesicles (GUVs):* GUVs were prepared using the electro swelling method[2] as described in.[3] Briefly, the lipid mixtures were prepared from stock solutions in EtOH. Subsequently, 100 nmol of POPC including 5 mol% of **5** or **6** were spotted onto custom-built titan chambers. These were placed on a heater plate at 50°C to facilitate solvent evaporation, and lipid-coated chambers were assembled using a spacer of Parafilm (Pechiney Plastic Packaging, Chicago, IL, USA) for insulation, subsequently put under high vacuum for at least 1 h in order to evaporate remaining traces of solvent. The electro swelling chamber was filled with 1 ml sucrose buffer (250 mM sucrose, 15 mM $NaN_3$, osmolarity of 280 mOsm/kg) and sealed with plasticine. An alternating electrical field of 10 Hz rising from 0.02 V to 1.1 V in the first 56 min was applied for 2.5 h at 55°C. To detach the vesicles from the slides the electrical field was changed to 4 Hz and 1.3 V and applied for 30 min.

*NMR measurements:* The respective molecules were mixed in the desired molar ratios in ethanol. After removing of the solvent, the samples were re-dissolved in cyclohexane and lyophilized overnight at high vacuum. Finally, the samples were hydrated with 50 wt% $H_2O$-buffer (10 mM Hepes, 100 mM NaCl, pH 7.4) and equilibrated by ten freeze-thaw cycles as well as gentle centrifugation. Static $^2$H NMR spectra were acquired on a Bruker (Bruker Biospin, Rheinstetten, Germany) DRX300 NMR spectrometer using a high-power probe with a 5-mm solenoid sample coil using the quadrupole echo pulse sequence.[4] The relaxation delay was 1 s, the delays between the 90° pulses (of ~3.2 µs) were 50 µs. Smoothed order parameter profiles were calculated [5], after the depaking the spectra.[6] The measurements were carried out at a temperature of 37°C.

*Measurement of fluorescence spectra:* Fluorescence spectra were recorded using an Aminco Bowman Series 2 spectrofluorometer (Urbana, IL, USA) with slit width for excitation and emission of 4 nm.

*Fluorescence lifetime measurements:* Fluorescence lifetime was measured as described.[7] Stock solutions of MLVs containing 2 mol% **5** or **6** were mixed with HBS and given into a fluorescence cuvette (final lipid concentration between 67 µM and 267 µM). The fluorescence lifetime was measured at room temperature with a FluoTime200 time-resolved spectrometer (PicoQuant GmbH, Berlin, Germany). A pulsed laser diode (LDH-P-C; PicoQuant GmbH, Berlin, Germany) of 405 nm and a pulse frequency of 8 MHz (125 ns pulse interval) were used as the excitation source. Individual photons were recorded on a time-correlated single photon counting setup with a time resolution of 33 ps. Emissions of **5** and **6** were recorded at 520 nm, and the spectral bandwidth was set to 16 nm. Data were acquired up to a level of 20.000 counts as defined by the maximum amplitude of the fluorescence lifetime decay kinetics. The instrument response function (IRF) was recorded using HEPES buffered salt solution at the excitation wavelength. Intensity decays were fitted globally (over at least seven data sets) using FluoFit software (PicoQuant GmbH, Berlin,Germany). A nonlinear least-squares iterative reconvolution fitting procedure was applied:

$$I(t) = \int_{-\infty}^{t} IRF(t')(\sum_{i=1}^{n} \alpha_i e^{-\frac{t-t'}{\tau_i}})\, dt' \quad (1)$$

with α being the amplitude of a lifetime component and τ being the corresponding lifetime. The lifetime decay kinetics were fitted with two exponential components. From these fittings, an average fluorescence lifetime ($\tau_{Av}$) was determined based on the intensity weighted ratio of two-time constants contributing to the biexponential kinetics according to:

$$\tau_{Av} = \frac{\sum_i \alpha_i \cdot \tau_i}{\sum_i \alpha_i} \quad (2)$$

where I(t) is the fluorescence intensity at time t and $\alpha_i$ is the preexponential factor representing the intensity of the time-resolved decay of the component with lifetime $\tau_i$.

*Measurement of UV spectra:* UV spectra were recorded at a V-630 spectrometer (JASCO, Pfungstadt, Germany). The emission spectra have been performed at a Fluoromax-4 spectrometer (Horiba Jobin Yvon, Bensheim, Germany) operated in the single-photon-counting mode. The fluorescence spectra were measured in a 90 degree angle to the excitation light. The samples were exited at their absorption maximum.

*Determination of the quantum yield:* The quantum yields have been measured at a photoluminescence quantum yield measurement system C9920 (HAMAMATSU Photonics, Herrsching, Germany) using an ULBRICHT sphere for an integral emission intensity collection. The samples were excited at their respective absorption maximum. The data were analyzed by the commercial software package provided by HAMAMATSU.

*Wide field epifluorescence microscopy:* For microscopy of GUVs, a Leica DMIRBE microscope equipped with an Andor IxonEM blue EMCCD camera operated at -75°C and a Lambda SC smart shutter (Sutter Instrument Company, USA) as illumination control was used. **5** and **6** were imaged with 100 × 1.3 NA oil immersion Fluotar objective (Leica Lasertechnik GmbH) using 360 nm (20 nm bandpass) excitation filter, 400 nm dichromatic mirror and 425 nm longpass emission filter. DilC18 was imaged using a standard red filter set (535 nm (50 nm bandpass) excitation filter, 565 nm dichromatic mirror, and 610 nm (75 nm) bandpass) emission filter. The resulting images were postprocessed by deconvolution using the ImageJ plugin DeconvolutionLab.[8] The Richard-Lucy algorithm was used with 30 iterations, and a theoretical point spread function (PSF) was used for deconvolution and generated using the Diffraction PSF 3D plugin in ImageJ (https://imagej.net/plugins/diffraction-psf-3d). Settings were chosen according to the used channel and camera specifications.

*Fluorescence life-time imaging microscopy (FLIM):* For FLIM measurements, an inverted FluoView 1000 laser scanning microscope (Olympus, Tokyo, Japan) equipped with a time-resolved LSM Upgrade kit (PicoQuant GmbH, Berlin, Germany) for time correlated single photon counting (TCSPC) was used. Images were obtained with a 60× water immersion objective (N.A. 1.2) at room temperature with a frame size of 512 × 512 pixels. The analogs **5** and **6** were excited with a pulsed 405 nm diode laser (pulse frequency 10 MHz; 4 µs/pixel), and fluorescence passed through 470/30 and 540/40 bandpass filters was detected using a τ single photon avalanche photodiode (τ-SPAD, PicoQuant GmbH). SPAD signals were processed with the TimeHarp 300 photon counting board and analyzed with the SymPhoTime 64 software (PicoQuant GmbH, Berlin, Germany) considering the instrument response function to allow consideration of short lifetime components with a high accuracy. FLIM images were acquired as described previously.[8]

*Fitting of the FLIM data*: Time-resolved photon-counts were summed up into a lifetime histogram within the SymPhoTime 64 software. The intensity distribution decay was analyzed by fitting using a nonlinear least squares iterative procedure as the sum of biexponential terms. For every single GUV, the intensity weighted average lifetime (τAv) was calculated according to:

$$\tau_{Av} = \frac{\sum_i \alpha_i \cdot \tau_i}{\sum_i \alpha_i} \qquad (2)$$

With α being the amplitude of a lifetime component and τ being the corresponding lifetime.

*Theoretical procedures*

*Molecular dynamics simulations:* Membrane bilayers with 70 POPC lipids and 30 sterols in each leaflet (200 lipids in total) were assembled with the Packmol program[9], using either cholesterol, **5**, or **6** as the sterol component. A pure POPC membrane containing 100 lipids in each leaflet was similarly assembled. The membranes also contained 10000 TIP3P water molecules[10], 22 $K^+$, and 22$Cl^-$ ions, corresponding to a salt concentration of about 0.15 M KCl. Force-field parameters for the POPC lipids were taken from lipid14. Parameters for the sterols were derived using the QFORCE program[11], based on r2scan-3c QM calculations[12] in the Orca program, version 5.0.3.[13] Charges were derived via fitting to the electrostatic potential, while bonded parameters were assigned using the QM hessian. Torsional parameters for flexible dihedrals were assigned by fitting to relaxed potential energy scans. Lennard-Jones parameters were taken from GAFF.[14]
All molecular dynamics simulations were performed using the Gromacs program, version 2022.3.[15] The assembled membranes were equilibrated in a three-step procedure consisting of minimization (5000 steps), equilibration in the NVT ensemble (200 ps), and equilibration in the NPT ensemble (2 ns). Long-range electrostatics were treated with the particle mesh Ewald method[16] with a short-range cutoff of 12 Å. A timestep of 2 fs was used, and bonds involving hydrogens were constrained using the LINCS algorithm.[17] The temperature was controlled by a Nose-Hoover thermostat[18] with a reference temperature of 298.15 K. In the NPT equilibration, a semi-isotropic Berendsen barostat was used[19], with a reference pressure of 1 bar. Production simulations were carried out with the same settings for 1000 ns but using a Parrinello-Rahman barostat.[20]
Order parameters of the POPC lipid tails were computed using the gmx order tool. Partial electron density profiles of selected membrane components were computed with cpptraj.[21] A hydrogen bond analysis was computed using vmd.[22] Sterol tilt angles and positions were extracted using the MDAnalysis Python library.[23]

Molecular Dynamics (MD) simulations of **5** and **6** solvated in water, ethanol, or mixtures of the two solvents were also carried out with the GROMACS program.[15] Initial cubic boxes with a side length of 12.7 Å containing 100 steroid molecules were assembled using the Packmol[9], corresponding to a concentration of about 80 mM. For each of the steroids, four solvent boxes were generated: a water box (50000 water molecules), an ethanol box (15000 ethanol molecules), a box with 50:50 volume ratio of the two solvents (7500 ethanol molecules, and 25000 water molecules) and a box with a 80:20 volume ratio of ethanol to water (12000 ethanol molecules, 10000 water molecules). The TIP3P model[10] was used for describing water molecules. Force field parameters for ethanol, **5,** and **6** were derived using the QFORCE package[11] as described above.

Each system was first minimized with the steepest descent algorithm with a force tolerance of 1000 kJ/mol and a maximum number of steps of 5000. After that, a 200 ps NVT simulation was performed at 298.15 K, with starting velocities generated by a Maxwell distribution at that temperature. A short 2 ns NPT simulation was carried out before the production run of each system. In all the MD runs, the timestep was 2 fs, and electrostatic interactions were evaluated with the Particle Mesh Ewald method[16], with a cut-off at 12 Å. Van der Waals interactions were cut off at 12 Å, with the Force-switch method between 10 Å and 12 Å. A Nose-Hoover thermostat and an isotropic Parrinello-Rahman barostat were employed for controlling, respectively, temperature and pressure at 298.15 K and 1 bar. The LINCS algorithm was used for constraining the bonds involving hydrogen atoms. All the production MD simulations were run for 500 ns. The different solvent environments were used for probing the aggregation capability of **5** and **6** as a function of the polarity of the solvent.

*Electronic structure calculations:* Structures of CTL, DHE, **5**, and **6** were optimized with CAMB3LYP-D3BJ/def2-TZVP[24] with the Orca program, version 5.0.3.[13] The optimized structures were used for subsequent DFT calculations[24a, 24d] of the excitation energies, oscillator strengths, and two- and three-photon absorption strengths using CAMB3LYP-D3BJ/def2-TZVP within the Dalton program, version 2022.[25] Two-photon cross-sections were converted to macroscopic units (GM), assuming a monochromatic light source with linear polarization and a Lorentzian broadening function with an FWHM of 0.1 eV. Natural transition orbitals were obtained with the Orca program. DFT calculations were also carried out on dimers of **5** and **6**. The molecules were oriented along their principal axes, and dimers were generated by rigid displacement along the *x*-axis, i.e., orthogonal to the plane of the ring systems, facing roughly the same direction as the C10-C19 bond vector. Excitation energies and associated oscillator strengths of the dimers were calculated as described above.

#### Supporting Figures and Tables

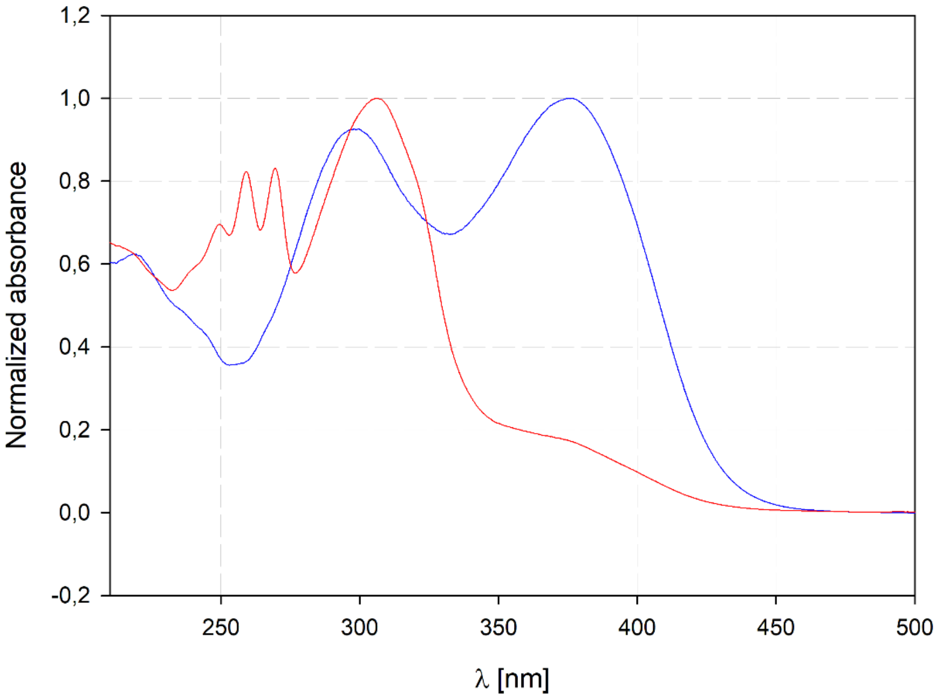


**Figure SI 1.** Normalized UV spectra of compounds **5** (blue line) and **6** (red line) in EtOH.

**Table SI 1.** Absorption maximum ($\lambda_{max}$), extinction coefficient (lg ε) and quantum yield ($\phi_F$) estimated for compounds **5** (blue line) and **6** (red line) in EtOH.

| Compound | $\lambda_{max}$ (lg ε) | $\Phi_F$ |
|---|---|---|
| **5** | 375 nm (4.1) | 25.6% |
| **6** | 306 nm (3.8) | 1.8% |

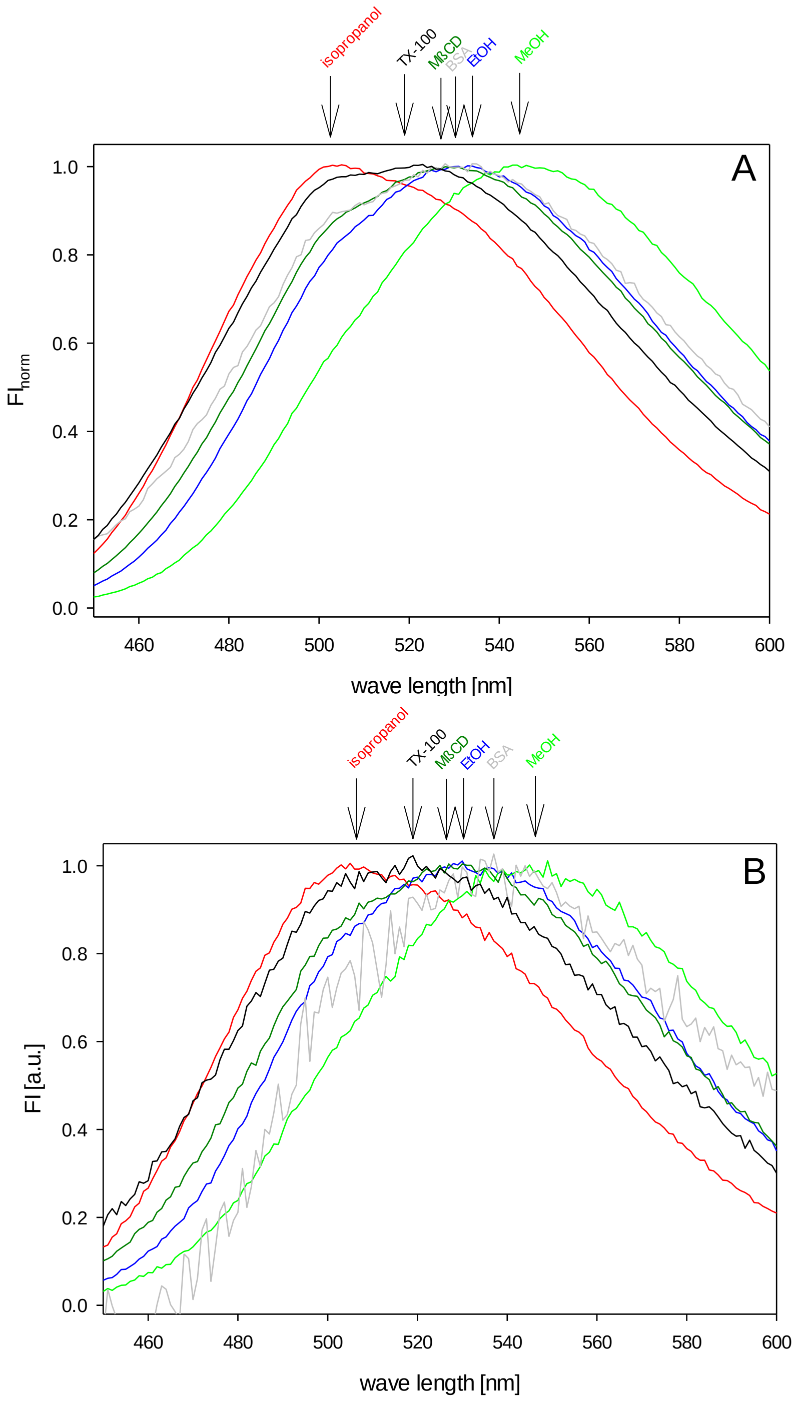


**Figure SI 2.** Fluorescence properties of **5** (A) and **6** (B) in different solvents. The fluorescent sterols were given from an ethanolic stock solution to the respective solvents or solutions, respectively (final concentration 10 µM): isopropanol (red line), ethanol (blue line), methanol (green line), Triton X - 100 (0.7%(v/v), (black line), methyl - β - cyclodextrin (MßCD) (1 mM, dark green line), and bovine serum albumin (BSA) (1%v/v), gray line). Fluorescence spectra ($\lambda_{ex}$ = 396 nm) were recorded at room temperature. From the spectra, the spectra of the solvents/solutions were subtracted. The spectra shown were normalized to the respective maximum fluorescence intensities.

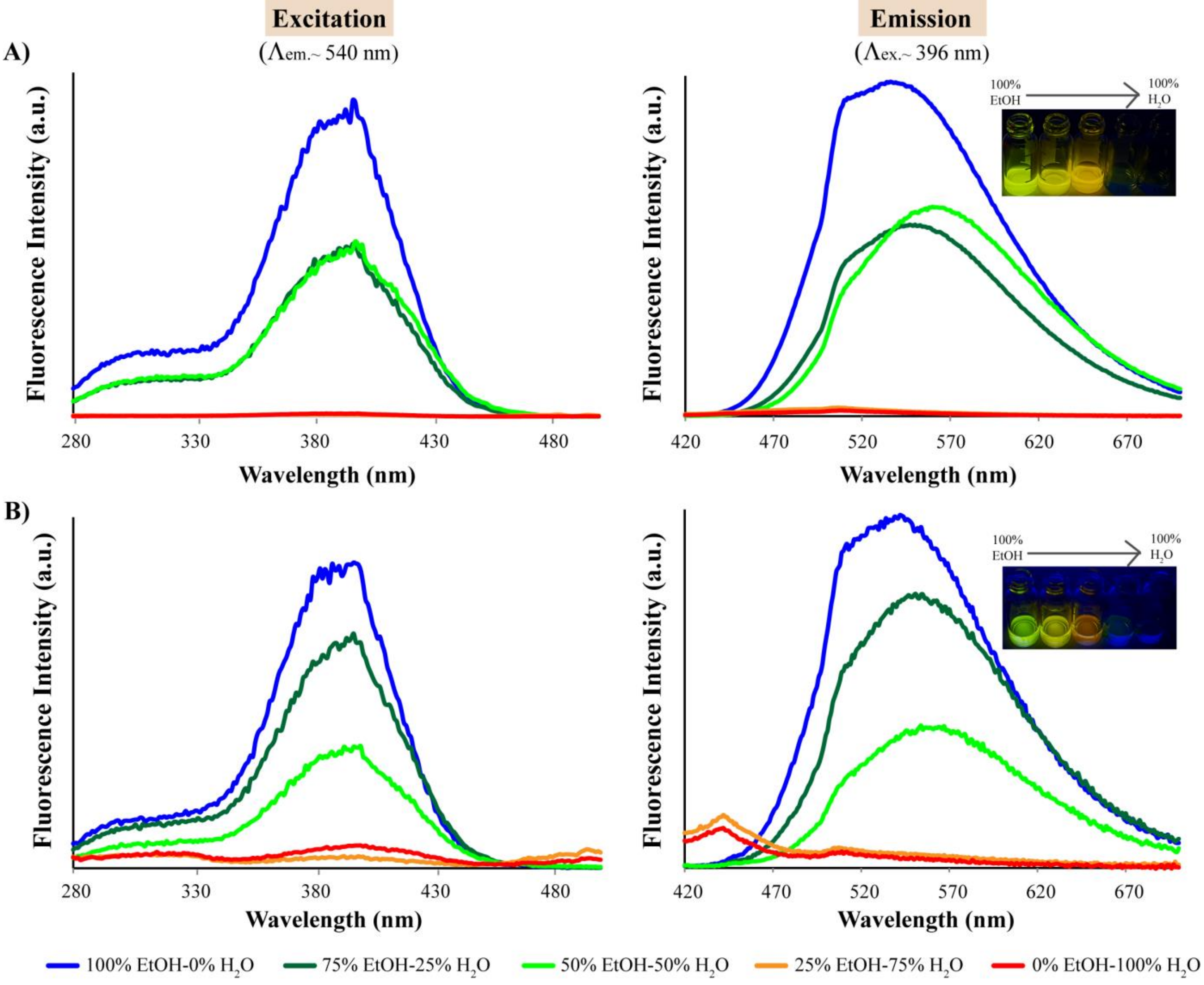


**Figure SI 3.** Excitation and emission spectra of **5** (A) and **6** (B) in water/ethanol mixtures.

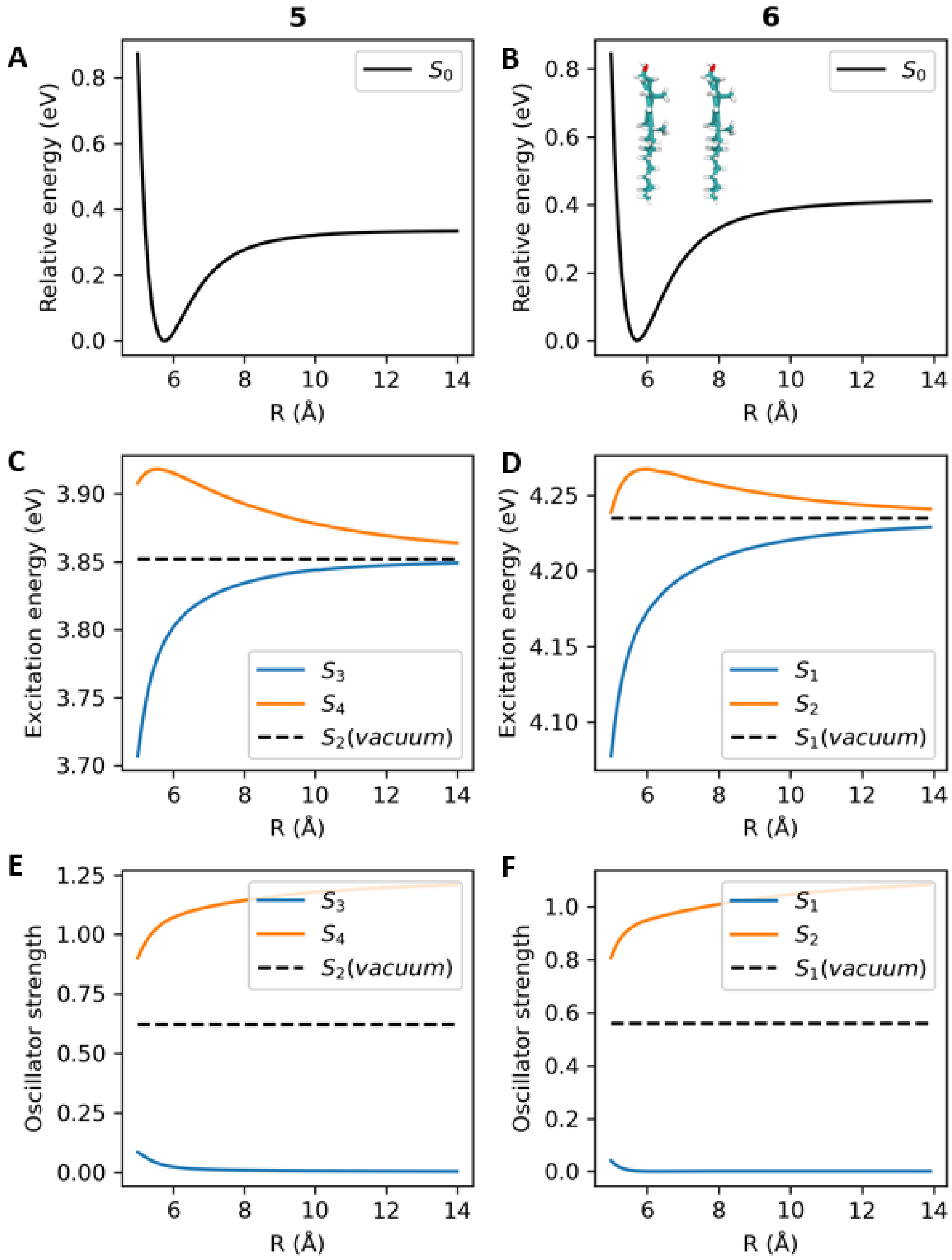


**Figure SI 4.** Calculation of exciton coupling in dimers of **5** and **6**. Calculation of exciton coupling in dimers of **5** and **6**. Dimers with a parallel orientation were generated by rigid displacement of the monomer geometries. Panels A, B show the ground-state interaction energy curves. Panels C, D show the excitation energies of the excitonically coupled states in the dimers. Panels E, F show the associated oscillator strengths of the transitions. For 5, the bright state of the isolated monomer is $S_2$; accordnigly, states $S_3$ and $S_4$ of the dimer are considered in the plots.

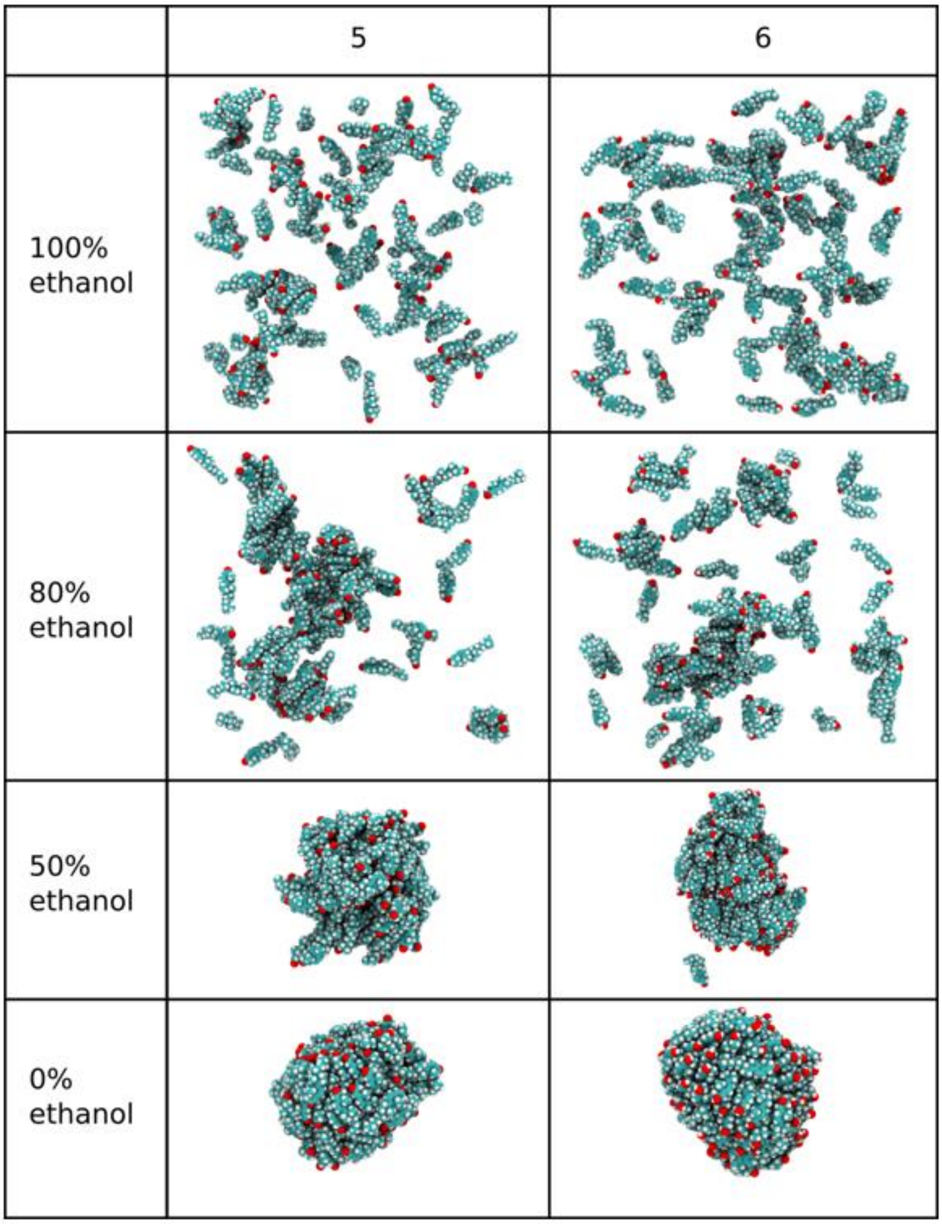


**Figure SI 5.** Molecular dynamics simulation of self-aggregation of **5** and **6** in water and water/ethanol mixtures.

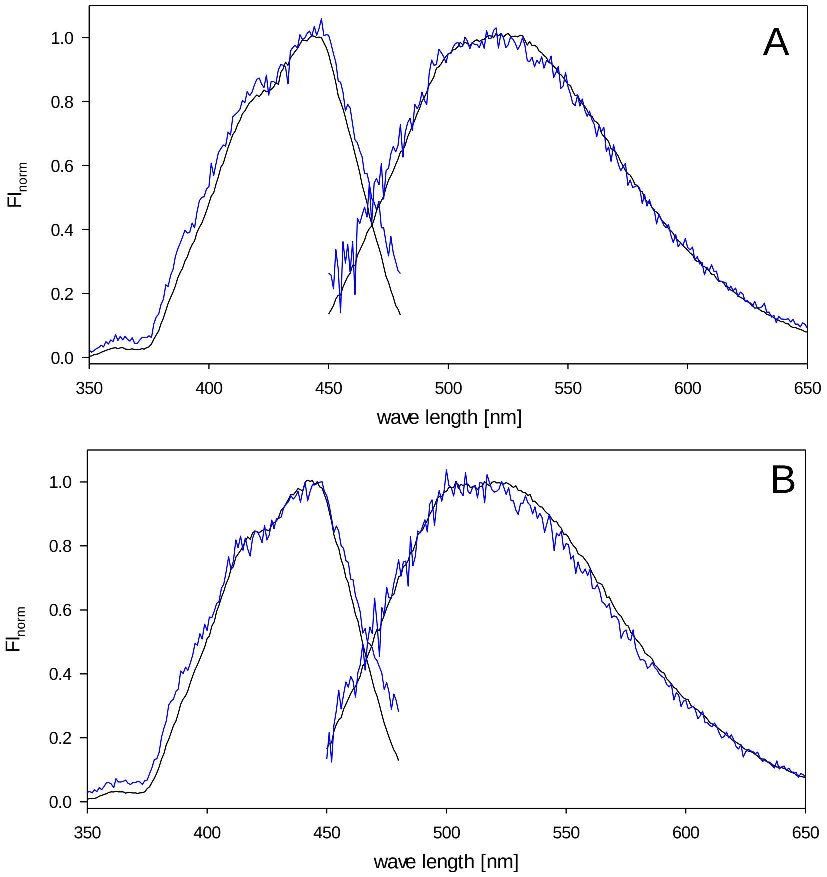


**Figure SI 6.** Fluorescence spectra of **5** and **6** in lipid membranes. Multilamellar vesicles consisting of 200 µM POPC (A) or 200 µM POPC/cholesterol (2:1) (B) and either 10 µM ML323 (black lines) or 10 µM ML349 (blue lines) were prepared. The fluorescence excitation spectra ($\lambda_{em}$ = 500 nm) and emission spectra ($\lambda_{ex}$ = 396 nm) of the vesicles were recorded at room temperature. The spectra were corrected for scattering by subtracting the spectra of respective unlabelled lipid vesicles and normalized to the respective maximum fluorescence intensities.

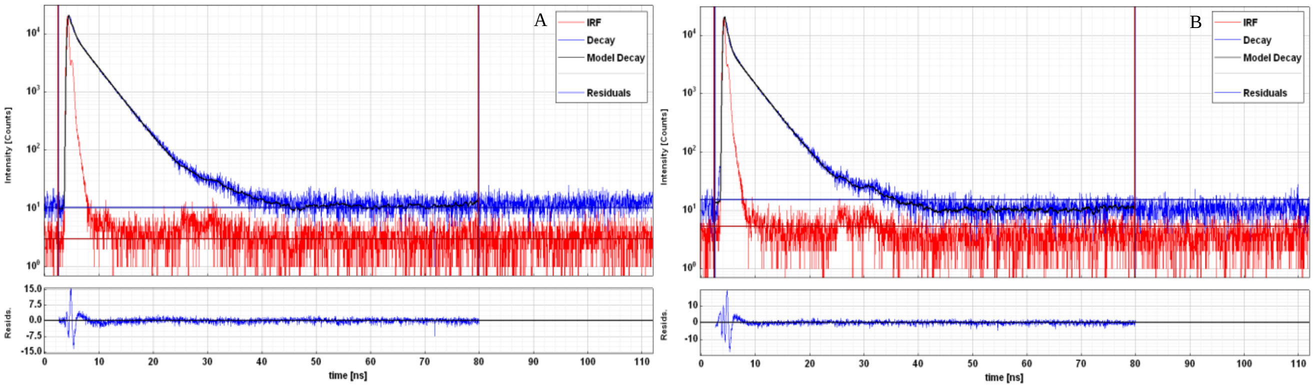


**Figure SI 7.** Fluorescence decay curves of **5** and **6** in lipid vesicles. Multilamellar vesicles containing POPC and 2 mol% of **5** (A) or **6** (B) were prepared and the decay curves were measured using a fluorescence lifetime spectrometer. The figure shows the decay curve of the respective analog, the IRF function as well as the model decay curve assuming a function with two exponential components and the respective residuals.

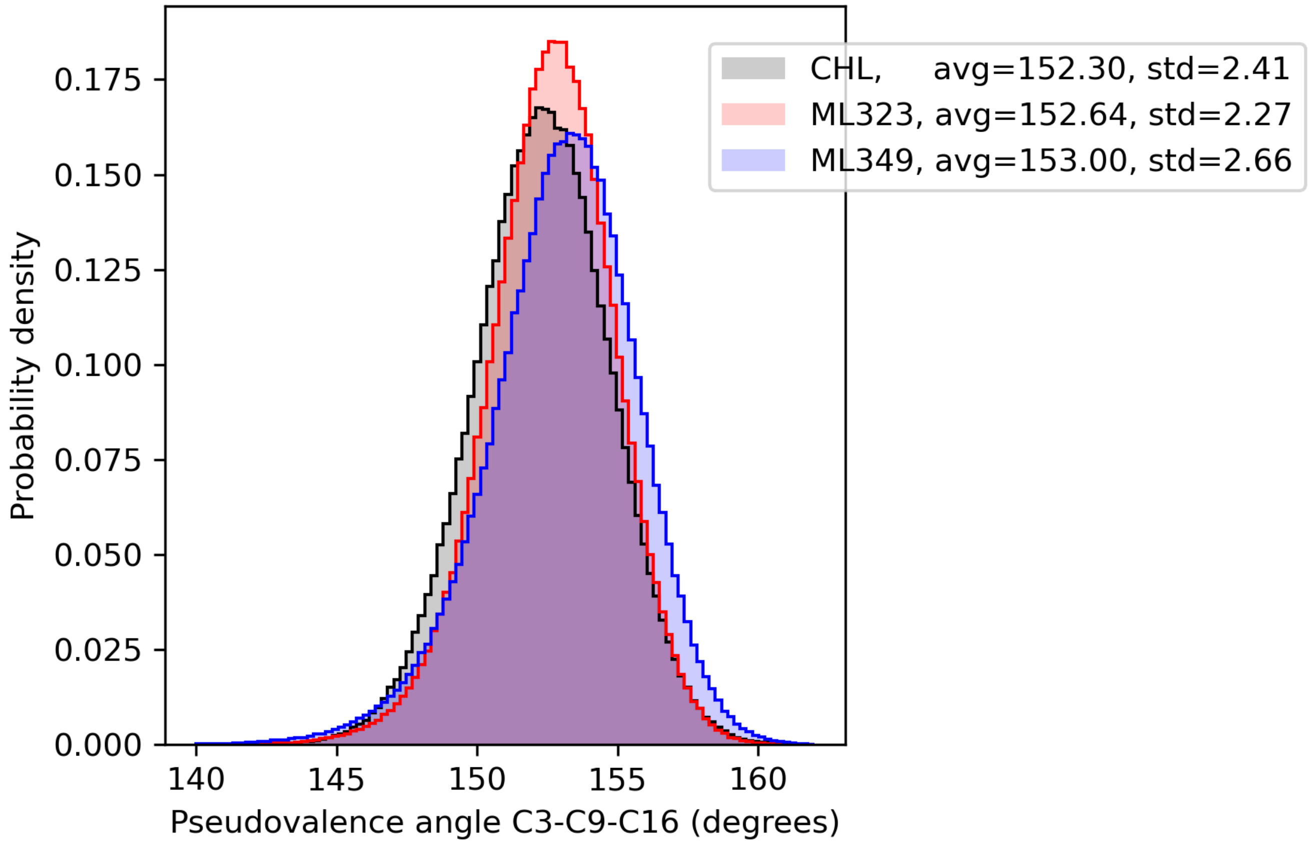


**Figure SI 8.** Ring flexibility assessed for **5** and **6** compared to cholesterol from molecular dynamics (MD) simulations. The distribution of pseudo angles between atoms 3, 9 and 16 of the steroid backbone was measured for cholesterol, **5** and **6** from the membrane simulations. The angle was sampled from 600000 snapshots of the respective trajectories.

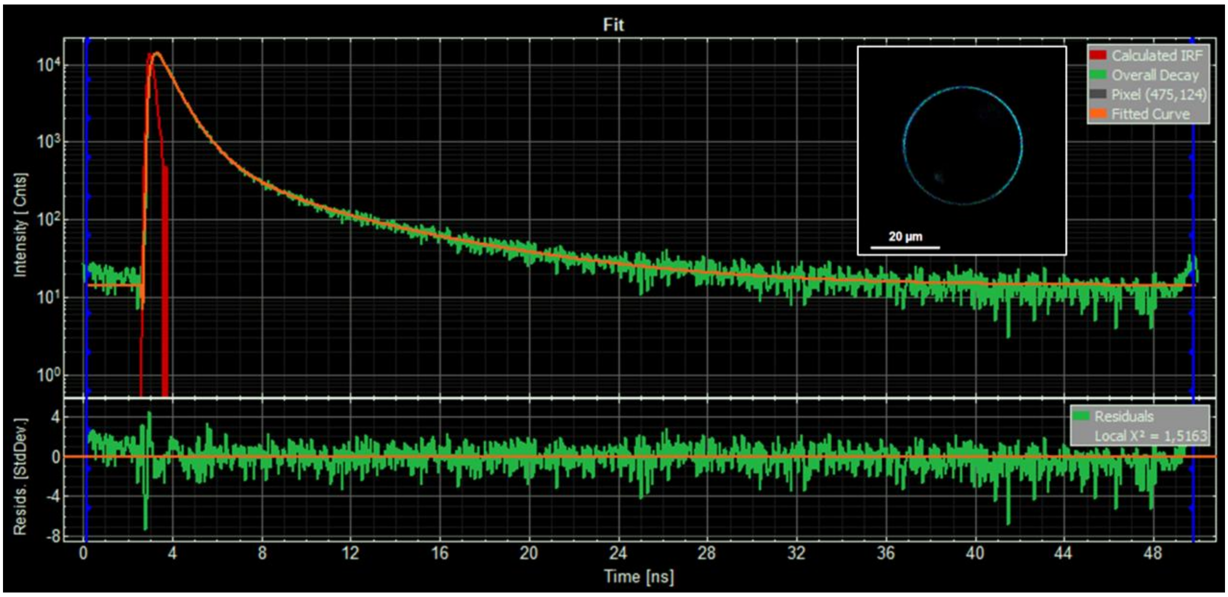


**Figure SI 9.** Fluorescence lifetime imaging of **5** in GUVs containing POPC and 5% **5**. GUVs were imaged at room temperature on an inverted FluoView 1000 laser scanning microscope (Olympus, Tokyo, Japan) equipped with a FLIM upgrade kit (PicoQuant GmbH, Berlin, Germany) using a water immersion objective (N.A. 1.2) with a frame size of 512 × 512 pixels. Time-resolved photon counts were summed up into a lifetime histogram within the SymPhoTime 64 software. The upper panel shows a typical decay curve of **5** (green), an imported IRF (red), and a tri-exponential curve fit (orange) with the residuals of the fit given in the lower panel. The inset shows a GUV as a color-coded fluorescence lifetime image.
